# The Cerebellar Engine: Multiscale Digital Brain Co-simulations Reveal How Cerebellar Spiking Architecture Shapes Cortical Coherence

**DOI:** 10.64898/2026.04.02.715849

**Authors:** Alice Geminiani, Jil Mona Meier, Dionysios Perdikis, Sabrine Ouertani, Claudia Casellato, Petra Ritter, Egidio D’Angelo

## Abstract

Cellular activities shape large-scale brain dynamics determining brain functioning and disease, yet the causal mechanisms across scales remain unclear. In particular, the cerebellum has been reported to modulate whole-brain dynamics during sensorimotor integration through unknown circuit interactions. To investigate the underlying mechanisms, we developed a novel multiscale digital brain simulator, in which a spiking neural network of the olivocerebellar microcircuit is embedded in a mean-field virtual mouse brain and wired using an atlas-based long-range connectome. Parameters were systematically tuned to match multiscale experimental data from primary sensory and motor cortices (S1 and M1) and cerebellum. We analyzed the role of cerebellar circuitry on sensorimotor integration by lesioning critical circuit connections *in silico*. Results suggested that Purkinje cell inhibition enhances the processing efficiency of the ‘cerebellar engine’ through decorrelation of cerebellar nuclei activity and that the pathway between mossy fibers and cerebellar nuclei is the specific pathway inside the microcircuit driving M1-S1 coherence. These results indicate a mechanistic link between cerebellar microcircuit and cortical sensorimotor processing. This novel framework opens new perspectives for the broader multiscale investigation of brain physiological and pathological states in relation to specific cellular and microcircuit properties.

## 1. Introduction

While animals interact with the external world, their brains continuously integrate information from multiple sensory modalities to control the planning, execution, and adaptation of movements, establishing a bidirectional link between motor commands and sensory feedback (Wolpert et al., 1995). This process, known as sensorimotor integration, is critical for flexible and skilled behavior across species (Sober & Sabes, 2005) and it is disrupted in motor disorders (Abbruzzese & Berardelli, 2003).

Sensorimotor integration relies on distributed cortical and subcortical circuits, including sensory and motor cortices, thalamus, cerebellum and basal ganglia (Romano et al., 2020), yet the exact contribution of each brain region remains unclear (Flanders, 2011). The “cerebellar engine” plays a crucial role in this process by integrating sensory and motor signals from multiple pathways (Bosman et al., 2011) and transforming multimodal sensory information streams into coordinated and precise motor adjustments, through synchronized neural activity across the primary sensory (S1) and motor (M1) cortices. At the network level, sensorimotor integration is characterized by gamma-band coherence. The cerebellum, thanks to its widespread connections with the forebrain (Palesi et al., 2020), is strategically positioned to boost cortical coherence during sensorimotor integration. The main question is then how the cerebellar circuitry contributes to cortical gamma coherence.

A prominent paradigm to study sensorimotor integration is rodent whisking, a naturalistic behavior where animals actively control rhythmic whisker movements and use tactile feedback from the whisker pad to infer environmental features and adjust ongoing motor output. In the complex brain circuit controlling whisking (Bosman et al., 2011), the cerebellum has been shown to be fundamental for movement amplification, adaptation and M1-S1 synchronization (Lindeman et al., 2021). Strikingly, experimental inactivation of the cerebellar nuclei (CN) – the output of the cerebellum – during free whisking significantly reduces M1-S1 gamma-band coherence (Popa et al., 2013). Cerebral cortex, thalamus and cerebellum can all generate gamma-band oscillations, and it has been proposed that the cerebellum may influence M1-S1 coherence via the cerebellothalamocortical pathway (Proville et al., 2014). However, the functional role of cerebellar microcircuit processing in shaping cortical dynamics during whisking is still poorly understood. Cerebellar Purkinje cells (PCs) encode position and velocity of whiskers (Chen et al., 2016) and can spontaneously generate gamma-band oscillations (De Zeeuw et al., 2008; Middleton et al., 2008), but whether and how their activity, together with other cerebellar microcircuit elements, is crucial for sensorimotor integration and the sensorimotor cortex coherence is difficult to isolate *in vivo* due to the technical challenges of performing highly specific manipulations during active behavior.

This experimental bottleneck opens a critical opportunity for computational multiscale modeling. While large-scale neuroinformatics platforms like The Virtual Mouse Brain (TVMB) excel at simulating macroscopic network dynamics using neural mass models of brain regions (Melozzi et al., 2017; Ritter et al., 2013), detailed spiking neural networks (SNNs) accurately capture local microcircuit computations (De Schepper et al., 2022; Geminiani et al., 2024). Recent proof-of-concept multiscale co-simulations have begun to bridge these macroscopic models of brain regions and spiking neural networks (Kusch et al., 2024; Meier et al., 2022; Schirner et al., 2023). However, these previous studies have focused on unidirectional effects and dynamics in the hippocampus or basal ganglia, not yet leveraging the unique feature of multiscale brain modeling to study the closed-loop effects between the scales. A framework that embeds a spiking cerebellum into a whole-brain network to investigate causal cross-scale interactions during a behavioral paradigm has so far been missing.

Here, we developed a novel multiscale model of the whole brain embedding a cerebellar SNN and used co-simulations to provide insights into the mechanisms of sensorimotor integration during a free-whisking paradigm. Following a multiscale fitting against multimodal datasets from literature, including the experimental study of (Popa et al., 2013), we tested the role of spiking cerebellar circuit architecture in sensorimotor integration *in silico* by lesioning connections inside the cerebellar microcircuit. We hypothesized that the cerebellum is not simply conveying signals into the M1-S1 loop, but it rather uses internal spike processing to control the coherence between M1 and S1 during sensorimotor integration. Our results highlight a potential specific microcircuit pathway driving M1-S1 coherence and provide evidence on the role of Purkinje cell inhibition in enhancing sensorimotor signal processing efficiency. Ultimately, this work provides a mechanistic hypothesis about how the cerebellum scales up microcircuit computations to orchestrate global brain dynamics, while also offering a widely reusable framework for investigating other complex cross-scale brain functions.

## 2. Results

### 2.1 Whole-brain co-simulations embedding a cerebellar spiking model: an *in silico* tool to investigate cerebellar mechanisms of sensorimotor integration

We developed a multiscale model of the rodent brain and simulated neural activity during free whisking to investigate how the cerebellum shapes cortical dynamics in sensorimotor integration. The connectivity of the whole-brain model was derived from the Allen Mouse Brain Connectivity Atlas (AMBCA) (Oh et al., 2014) (Fig. 1A). Cortical brain regions were modelled as cortico-thalamic Wilson-Cowan neural masses (Griffiths et al., 2020) and subcortical ones as a modified version of the same neural mass model without the loop to the specific thalamus (Fig. 1C). An SNN model was used for the cerebellar cortex and its main whisking input region (the principal nucleus of the trigeminus), the CN and the inferior olivary complex (IO) (De Schepper et al., 2022; Geminiani et al., 2024) (Fig. 1D). The cerebellar SNN receives sensory inputs – including whisking signals from the trigeminal nuclei – via mossy fibers, which reach the CN (output of the cerebellum) through two main pathways: a direct excitatory projection from mossy fiber collaterals (direct *nuclear* pathway) and an indirect inhibitory projection from PCs, which conveys the result of cerebellar cortical processing of mossy fiber signals (indirect *cortical* pathway) (Fig. 1D). Finally, whisker neural mass nodes allowed for whisking to be simulated with a closed loop between the cortex and the periphery via the most important subcortical regions involved in transferring the whisking-related signals, i.e., a minimal whisking pathway, with the involved connections amplified by a pathway gain to simulate the task (Fig. 1B). The neural mass nodes were simulated with TVMB (Melozzi et al., 2017), an extension of The Virtual Brain (Ritter et al., 2013; Sanz-Leon et al., 2013; Schirner et al., 2022), while the SNN was implemented in NEST.

**Figure 1.**
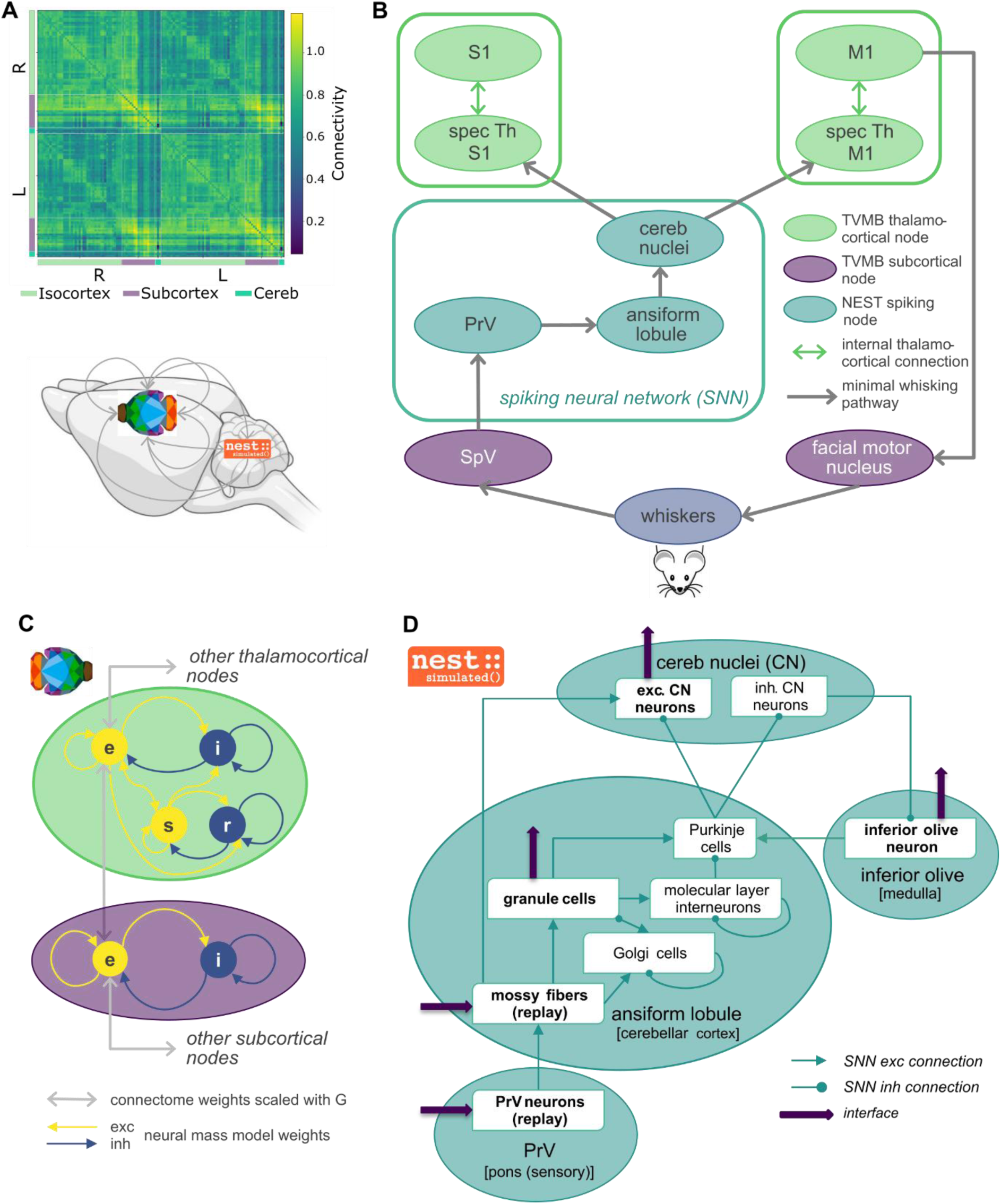
Overview of the whole-brain model. **(A)** Whole-brain connectivity matrix (without specific thalami and whiskers; diagonal values are zero but are set to the average matrix value for visualization) derived from the Allen Mouse Brain Connectivity Atlas. Mouse brain icon from bioRender.com. **(B)** Schematic of the minimal whisking pathway, along which connections are scaled with the pathway gain. The network includes thalamocortical (TC) and subcortical nodes implemented in The Virtual Mouse Brain (TVMB) and spiking nodes implemented in the NEural Simulation Tool (NEST). **(C)** Schematic of TVMB nodes, including thalamocortical nodes (in green) with four state variables (e: excitatory neural activity, i: inhibitory neural activity, s: excitatory relay neural activity, r: inhibitory reticular neural activity) and subcortical nodes (in purple) with excitatory and inhibitory neural activity state variables. **(D)** Spiking neural network implemented in NEST, with neuronal populations and internal connections. The network includes the ansiform lobule node (from the cerebellar cortex) with four neuronal populations, the cerebellar nuclei (CN) node with two neuronal populations, the inferior olive node (from the medulla) with one neuronal population. The interface points (input and output) with TVMB are indicated by red arrows. The overall sensory input from TVMB is conveyed to the ansiform lobule and the CN through mossy fibers. These fibers also receive the whisking input from replay neurons in the principal sensory nucleus of trigeminus that in turn receives the input from TVMB through an interface point. Output from the spiking network comes from granule cells of the ansiform lobule (to the rest of the cerebellar cortex through parallel fibers), excitatory neurons of the CN and inferior olive neurons. Fibers/neural populations at the interface are marked in bold. S1: primary somatosensory area, barrel field; M1: primary motor area; spec Th: specific thalamus; cereb: cerebellum/cerebellar; PrV: principal sensory nucleus of the trigeminus; SpV: spinal nucleus of the trigeminus; SNN: spiking neural network; exc: excitatory; inh: inhibitory; conn.: connection; G: global coupling scaling.

### 2.2 The model captures experimental data across scales

Free parameters in the model were systematically tuned against multiscale datasets from rodent whisking experiments, including local field potentials in M1 and S1, and single-unit electrophysiology recordings in the cerebellum. The resulting model reproduced experimental phenomena across scales, from neural field signals to single-neuron spiking activity, both in measures used as targets during fitting and in other relevant measures for the biological plausibility of simulations (Fig. 2). After systematic model parameter tuning against experimental data (<u>Par. 4.2.1</u>), PSDs of M1 and S1 from TVMB-only simulations were similar to the PSDs of local field potentials from rodent recordings in the same brain regions (Popa et al., 2013) (Fig. 2A-B), with average values in theta- and gamma-bands close to the target (Fig. 2C). Similar M1 and S1 PSDs, with comparable values of theta- and gamma-band average power, were obtained in TVMB-NEST co-simulations (Supplementary Fig. S1), demonstrating robustness of fitting when increasing simulation complexity.

**Figure 2.**
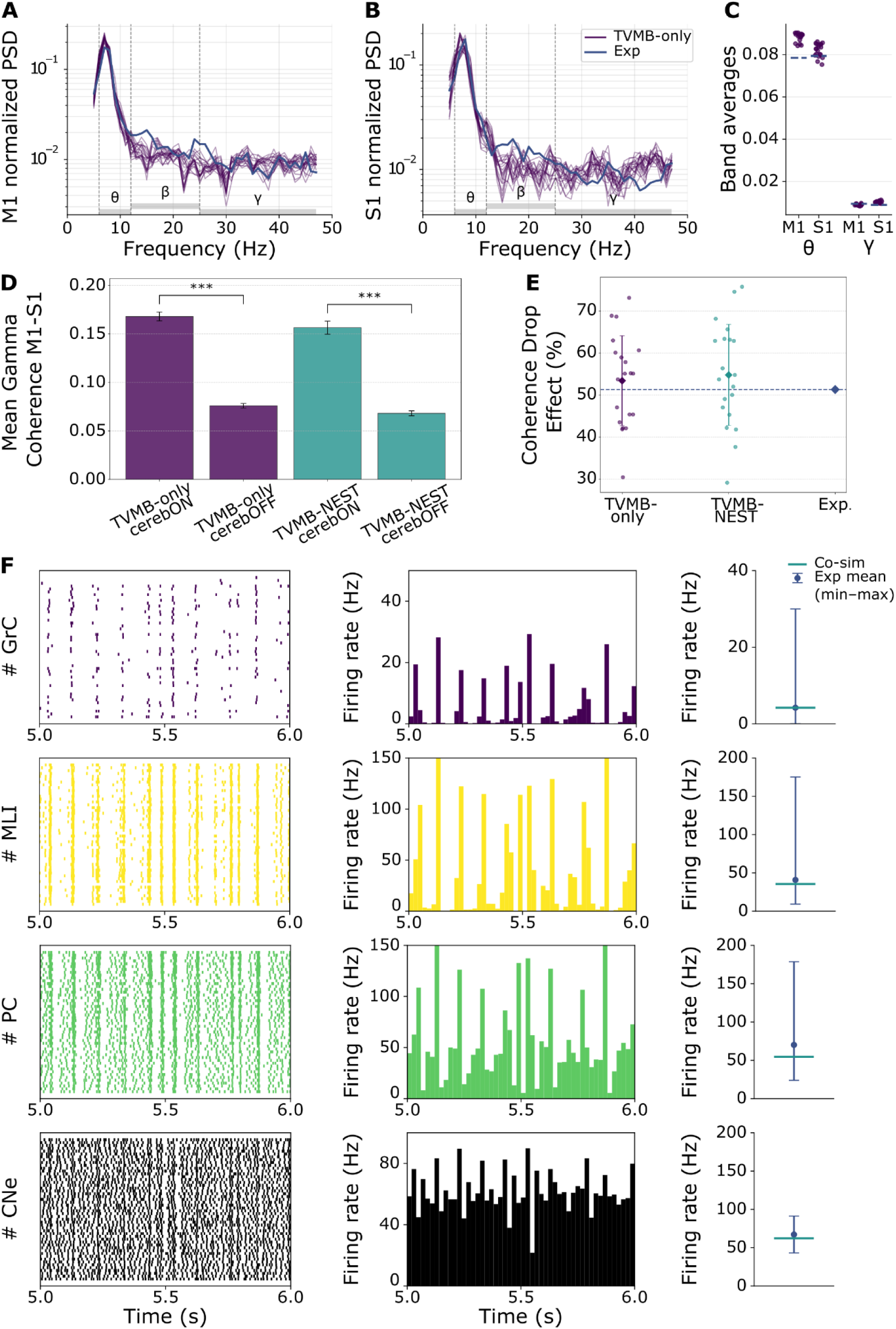
Functional validity of the model across scales. **(A)** PSD of M1 for 10 TVMB-only simulation runs and both hemispheres (n=20), in purple, compared to the target experimental PSD from (Popa et al., 2013), in blue. **(B)** Same for S1. **(C)** Average power in theta and gamma bands. **(D)** M1-S1 coherence drop in gamma band (25 – 60Hz) when inactivating the cerebellum, in TVMB-only simulations and TVMB-NEST co-simulations. Bar plots of mean gamma coherence across 10 simulation runs, with standard deviations as error bars, for TVMB-only simulations (purple) and TVMB-NEST co-simulations (green). **\*\*\*** indicates a significant difference (two-sided t-test, p < 0.0001). **(E)** Coherence drop effect as percentage of the normalized difference (cerebON – cerebOFF) / cerebON in TVBM-only and TVMB-NEST co-simulations, displayed as means and standard deviation (over the pooled hemispheric values), where dots represent single simulation runs of each hemisphere. The rightmost average value displays the experimental data from (Popa et al., 2013), where mean gamma-band coherence drops between M1 and S1 were similarly calculated for comparison. Both TVMB-only (mean: 53.4%) and co-simulations (mean: 54.8%) have mean coherence drop values close to the experimental data (51.3%). **(F)** Raster plots from 50 sampled neurons (left) and 20ms-bin spike population histograms (center) for the main cell types in the cerebellar SNN of the right hemisphere (granule cells (GrCs), molecular layer interneurons (MLIs), Purkinje cell (PC), excitatory neurons in the cerebellar nuclei (CNe)) during a 1s interval of an example 10s simulation of whisking. The plots on the right report the mean across the population for 10 simulation runs (light green line) compared to experimental mean (blue dot) and range (min to max, blue error bar) from (Chen et al., 2017) for GrCs and MLIs, (Chen et al., 2016) for PC and (van der Heijden et al., 2022) for CNe. S1: primary somatosensory area, barrel field; M1: primary motor area; PSD: power spectral density; Exp.: experimental data; TVMB: The Virtual Mouse Brain; NEST: NEural Simulation Tool; Co-sim: TVMB-NEST co-simulation results.

The pathway gain parameters were optimized using TVMB-only simulations, against the experimental M1-S1 gamma-band coherence drop when inactivating CN (Popa et al., 2013). Specifically, we measured M1-S1 gamma-band coherence within each brain hemisphere and its drop between *cerebON* (active cerebellum) and *cerebOFF* (virtually inactivated cerebellum) conditions. Both TVMB-only and TVMB-NEST co-simulations reproduced the significant drop in M1-S1 gamma-band coherence during whisking when inactivating the cerebellum (Fig. 2D-E). Notably, the optimized pathway gains also facilitated the emergent propagation of the whisking signal: the regions involved exhibited theta-band oscillations representing whisker movement (e.g., Fig. 2F and 3A), although these oscillations were not an explicit fitting target.

**Figure 3.**
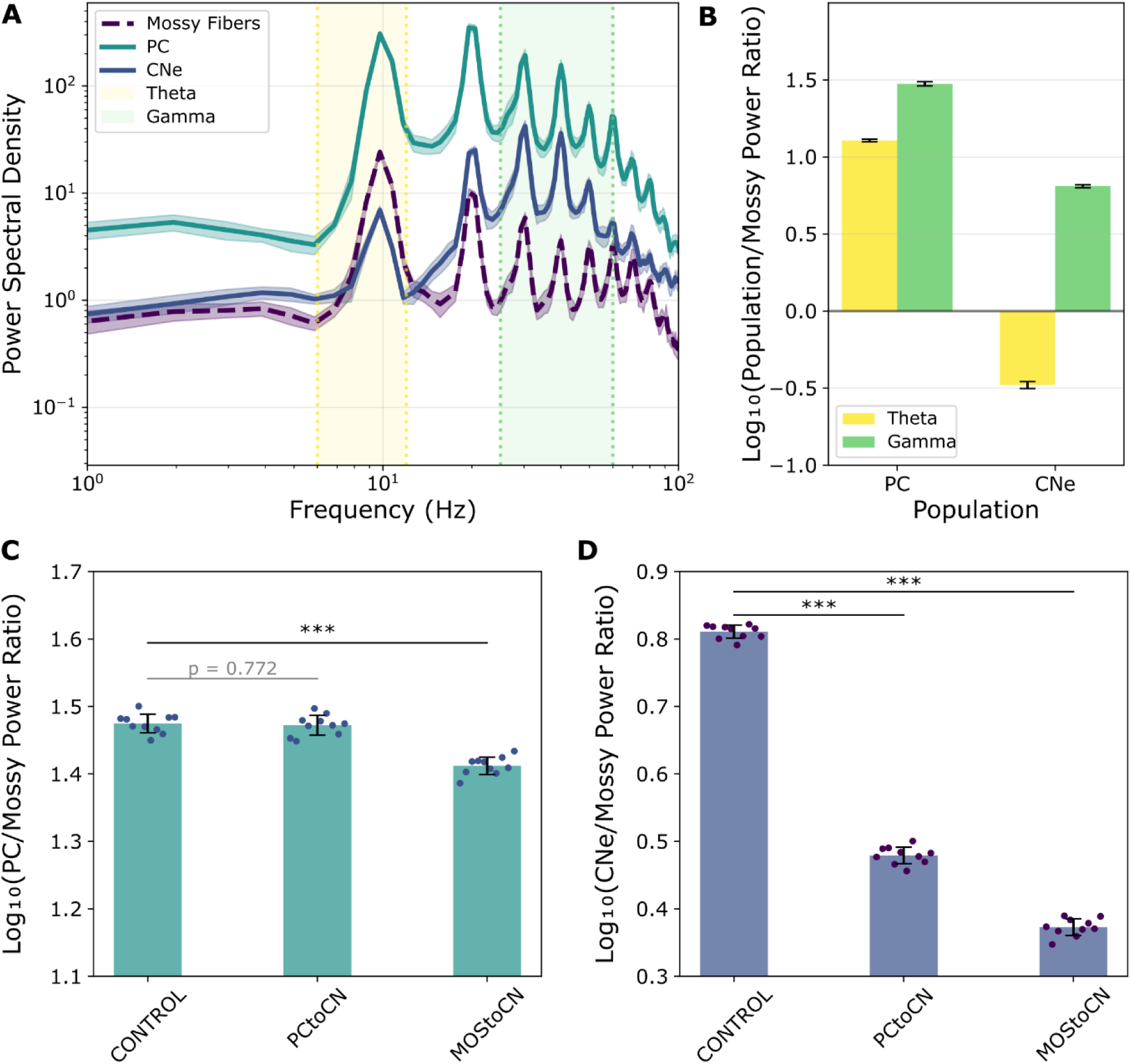
Oscillatory power inside the cerebellar circuit. **(A)** Power spectral density of mossy fibers (cerebellar input), Purkinje cells (PC, cerebellar cortex output) and excitatory cerebellar nuclei neurons (CNe, cerebellar output) in CONTROL. Lines and shaded areas represent mean and standard deviation of 10 simulation runs, each averaged over both hemispheres. **(B)** Bar plot of the average relative increase of gamma- and theta-band power in the two populations of interest (PC and CNe) with respect to mossy fibers, expressed as log_10_ of the power ratio. Error bars display standard deviations across 10 simulations, each averaged over both hemispheres. **(C-D)** Relative change of power in the two populations of interest – Purkinje cells, PC (C) and excitatory cerebellar nuclei neurons, CNe (D) – with respect to mossy fibers measured as gamma-band power ratio in CONTROL and in virtual lesion conditions – PCtoCN and MOStoCN. (n = 10 independent samples, averaged over both hemispheres, mean ± STD, ***p < 0.001; two-sided unpaired t-test). PCtoCN: inactivating the indirect inhibitory projection from PC to CNe; MOStoCN: inactivating the direct excitatory projection from mossy fibers to CNe; CNe: excitatory cerebellar nuclei neurons.

Finally, using single-unit electrophysiology data during whisking (Chen et al., 2017), the inter-scale interface parameters for TVMB-NEST co-simulations were tuned applying grid search to obtain physiological firing rates of granule cells in the cerebellar SNN (Supplementary Note S1.2 and Supplementary Figure S2). The results were robust for different configurations of parameters, which correspond to a range of values of *G*, the global coupling of the large-scale connectivity (Supplementary Fig. S2 and S3). In co-simulations, the whisking signal was effectively transmitted from the principal sensory nucleus of the trigeminus (PrV) to the cerebellar SNN via the tuned inter-scale interface: all major cerebellar neuronal populations exhibited whisking-related theta-band oscillations (Fig. 2F-left and middle column), which would be absent without a theta-band oscillating input (Geminiani, Pedrocchi, et al., 2019), and average firing rates within physiological ranges (Fig. 2F-right column). Although only the granule cell firing rate was used to fit the input interface parameters, all other cerebellar populations also displayed physiologically realistic firing rates, as measured in rodent single-unit recordings during free whisking (Chen et al., 2016, 2017; van der Heijden et al., 2022). Taken together, these results support the functional validity of the SNN within the whole-brain model and its generalization from resting-state and eyeblink conditioning (Geminiani et al., 2024; Geminiani, Pedrocchi, et al., 2019) to whisking.

Experiments show that PC activity exhibits gamma-band oscillations, (Lindeman et al., 2021; Middleton et al., 2008), and they play a crucial role in shaping gamma-band oscillations in the CN (Shin & De Schutter, 2006), in synergy with mossy fiber synaptic inputs (Y. Wu & Raman, 2017). It has been hypothesized that gamma-band oscillations could in turn be relevant for M1-S1 gamma-band coherence (De Zeeuw et al., 2008), but experimental evidence is still inconclusive (McAfee et al., 2022). To test this hypothesis, we first evaluated whether the SNN, which was not tuned against any oscillatory property, could reproduce the experimental gamma-band oscillations (Middleton et al., 2008; Person & Raman, 2011). We measured the power of PCs and CNe, relative to mossy fibers, to normalize for any contribution of incoming oscillations from outside the cerebellum (Fig. 3B). Gamma and theta power of PCs increased with respect to mossy fibers’ power, while power increased in the gamma- and decreased in the theta-band for CNe relative to the mossy fibers (Fig. 3A-B).

### 2.3 Virtual lesions in the cerebellar spiking neural network reveal the role of distinct cerebellar circuit elements in regulating cerebellar and cortical gamma-band dynamics

We exploited the higher level of granularity in the cerebellar SNN model to study the role of different cerebellar mechanisms on cortical coherence during the whisking task. In TVMB-NEST co-simulations, we applied localized virtual lesions to the SNN and compared the results to those obtained in non-lesioned conditions (*CONTROL*). To isolate the contribution of each connection to population dynamics independently of changes in mean activity (Heinzle et al., 2007), we adjusted single-neuron intrinsic parameters of the target populations to preserve their average firing rates and normalized the cerebellar output signal.

#### 2.3.1 Cerebellar microcircuit contributions to gamma-band power

To investigate the contribution of PC and mossy fibers in shaping the cerebellar output (Y. Wu & Raman, 2017), we ran TVBM-NEST co-simulations, while independently inactivating input connections to the CNe, i.e., the excitatory connections from mossy fibers (*MOStoCN*) and the inhibitory ones from PCs (*PCtoCN*). The PC relative to mossy fiber power was unchanged with *PCtoCN*, as expected given that the pathway from mossy fibers to PC remained intact, whereas it was significantly reduced with *MOStoCN* (Fig. 3C). Instead, relative CNe gamma-band power was significantly decreased with both lesions (Fig. 3D). Whereas the mossy fiber signal might change, the gamma-band power of mossy fibers did not change significantly in both micro-lesions and the absolute PC and CNe gamma-band power changed in the same directions as the relative one (Supplementary Fig. S4).

#### 2.3.2 Cerebellar microcircuit contributions to M1-S1 gamma-band coherence

We explored how M1-S1 gamma-band coherence during whisking is affected by processing within the cerebellum, i.e., isolating possible effects due to the connections between sensorimotor cortex and cerebellum. To that end, we asked how the cortical and nuclear cerebellar pathways influence gamma-band coherence, not only in the sensorimotor cortex, but also between the PrV, i.e., the input region to the cerebellum, and the CN, the output cerebellar region towards the sensorimotor thalamocortical loops. We compared those two coherence effects for the *CONTROL* condition and the two lesions disrupting the direct and indirect pathways in the cerebellum (*MOStoCN* and *PCtoCN*, respectively).

Both lesions had distinct effects on cerebellar input-output and M1-S1 gamma-band coherence (Fig. 4). Removal of excitatory connections from mossy fibers (*MOStoCN*) decreased cerebellar input-output coherence and M1-S1 coherence to a level like the one with inactivation of the cerebellum (Fig. 2D), whereas removal of inhibitory connections from PCs (*PCtoCN*) increased both coherences. When examining gamma-to-theta coherence ratio, both lesions reduced cerebellar input-output and M1-S1 values (Fig. 4C-D).

**Figure 4.**
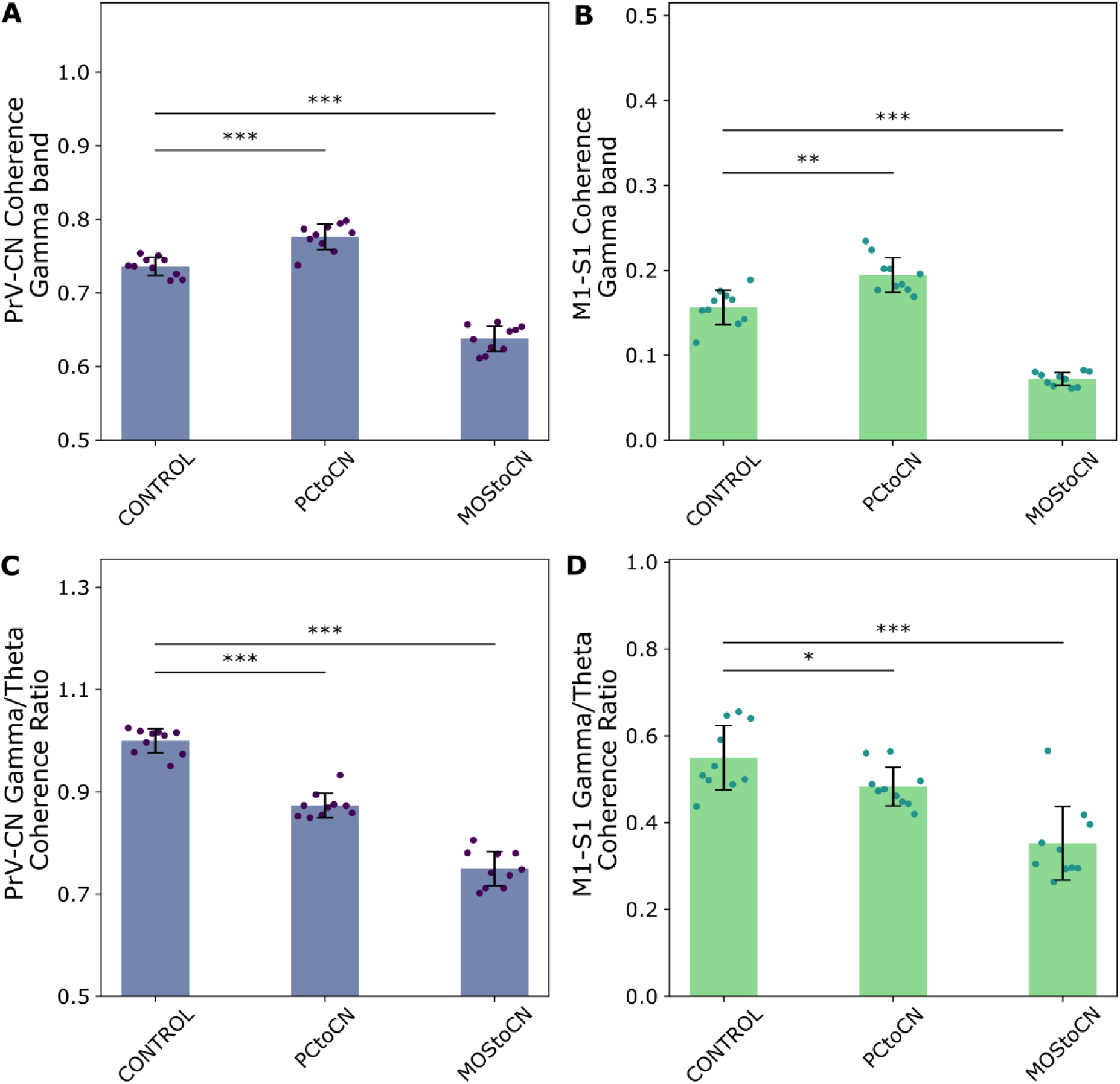
Gamma-band coherence with input to CN lesions. **(A-B)** Gamma-band and **(C-D)** gamma-to-theta coherence for PrV-CN (cerebellar input-output) and M1-S1 in CONTROL and with separate ablations of synaptic inputs to the CN. Bar plots and error bars respectively show the mean and standard deviation over 10 simulations, each one averaged across both hemispheres, with comparisons between CONTROL and each lesion condition (*p < 0.05, **p < 0.01, ***p < 0.001; independent-samples two-sided unpaired t-test, FDR-corrected over all lesion conditions). PCtoCN: inactivating the indirect inhibitory projection from PC to CNe; MOStoCN: inactivating the direct excitatory projection from mossy fibers to CNe; PrV: principal sensory nucleus of the trigeminus; CN: cerebellar nuclei; S1: primary somatosensory area, barrel field; M1: primary motor area; CNe: excitatory cerebellar nuclei neurons.

To assess the role of cerebellar output’s baseline and amplitude changes to the coherence effect, we ran simulations also without normalization of the cerebellar output. Gamma-band coherence trends across conditions remained evident in these simulations (Supplementary Fig. S5). Therefore, observed changes in coherence were independent of the lesion-induced changes in average cerebellar output activity - increased for inhibitory *PCtoCN* and decreased for excitatory *MOStoCN*. However, the effect on gamma-to-theta coherence ratio was masked by changes in the CN output activity in non-normalized simulations (Supplementary Fig. S5CD).

Recent experimental evidence points out that PCs can generate gamma-band oscillations supposedly thanks to molecular layer inhibition (De Zeeuw et al., 2008; Middleton et al., 2008). However, it is not known whether MLI inhibition also impacts cerebellar gamma-band coherence and cortical gamma-band coherence indirectly through the CN output. To address this question in our model, we ran co-simulations in two SNN lesion conditions: (i) directly inactivating MLI to PC inhibition (*MLItoPC*) and (ii) inactivating self-inhibition in the molecular layer (*MLItoMLI*) to analyze its indirect effect on PC dynamics. Our simulations show that MLI inhibition regulates PC gamma-band dynamics without affecting M1-S1 coherence (Supplementary Note S2).

#### 2.3.3 Cerebellar circuit elements differentially contribute to gamma-band power and coherence

When considering the effects of all lesions on gamma-band power and coherence together (Table 1), our results show that the lesions impact these measures differently. Regarding gamma power, it generally drops in the cerebellum for all lesions except for *MLItoMLI*, this effect being more prominent for *MOStoCN* and *PCtoCN*. Coherence, though, drops mainly for the direct pathway lesion (*MOStoCN*). For the indirect pathway lesions, coherence drops only for *MLItoPC,* to a lesser extent and only within cerebellum, whereas it increases for *PCtoCN* and it is unaffected for *MLItoMLI*. On the other hand, changes in cerebellar input-output coherence (both computed between the whisking input region – PrV – and the CN region (Table 1) and between mossy fibers and CNe activities inside the SNN (Supplementary Fig. S7)) and M1-S1 coherence occurred in the same direction, with the only difference that in some cases they did not transfer to the sensorimotor cortex.

**Table 1.**
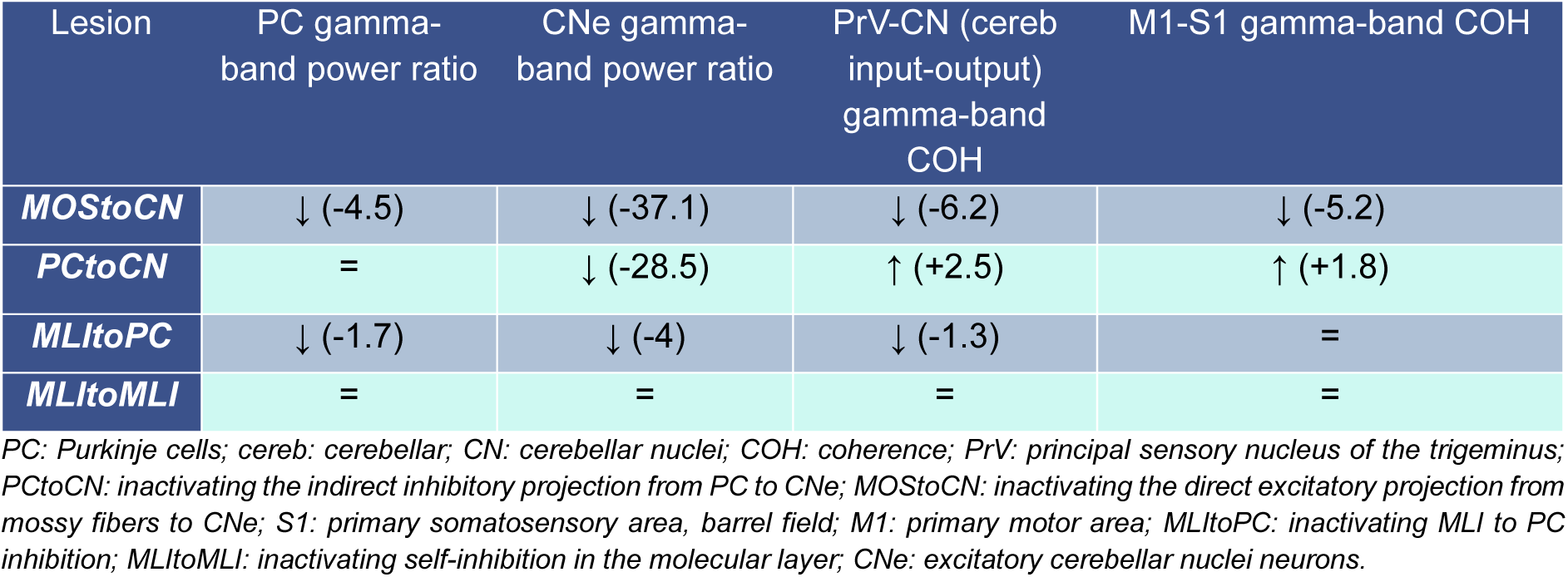
Oscillatory changes with cerebellar microcircuit lesions. Summary of effects of the lesions on PC and CNe gamma-band power relative to mossy fibers, on cerebellar input-output coherence, and M1-S1 coherence in gamma band. ↓, ↑ and = indicate significantly lower, significantly higher and non-significantly different, respectively, compared to CONTROL. Values in brackets are the Cohen’s distances quantifying the effect size of each significantly different comparison.

| Lesion | PC gamma-band power ratio | CNe gamma-band power ratio | PrV-CN (cereb input-output) gamma-band COH | M1-S1 gamma-band COH |
| --- | --- | --- | --- | --- |
| *MOSToCN* | ↓ (-4.5) | ↓ (-37.1) | ↓ (-6.2) | ↓ (-5.2) |
| *PCtoCN* | = | ↓ (-28.5) | ↑ (+2.5) | ↑ (+1.8) |
| *MLItoPC* | ↓ (-1.7) | ↓ (-4) | ↓ (-1.3) | = |
| *MLItoMLI* | = | = | = | = |

PC: Purkinje cells; cereb: cerebellar; CN: cerebellar nuclei; COH: coherence; PrV: principal sensory nucleus of the trigeminus; *PCtoCN*: inactivating the indirect inhibitory projection from PC to CNe; *MOSToCN*: inactivating the direct excitatory projection from mossy fibers to CNe; S1: primary somatosensory area, barrel field; M1: primary motor area; *MLItoPC*: inactivating MLI to PC inhibition; *MLItoMLI*: inactivating self-inhibition in the molecular layer; CNe: excitatory cerebellar nuclei neurons.

We computed the cross-spectral density (CSD) between mossy fibers and CNe (*COH_xy_* = |*P_xy_* | ⁄(*P_x_ P_y_*) with |*P_xy_*| being the magnitude of the CSD). This quantity provides a measure of global changes – power and coherence combined – between mossy fibers and CNe oscillations that are relevant for brain oscillations and behavior downstream to the cerebellum during sensorimotor integration (Lindeman et al., 2021). Both lesions to CN inputs decreased CSD magnitude in the gamma band (Fig. 5).

**Figure 5.**
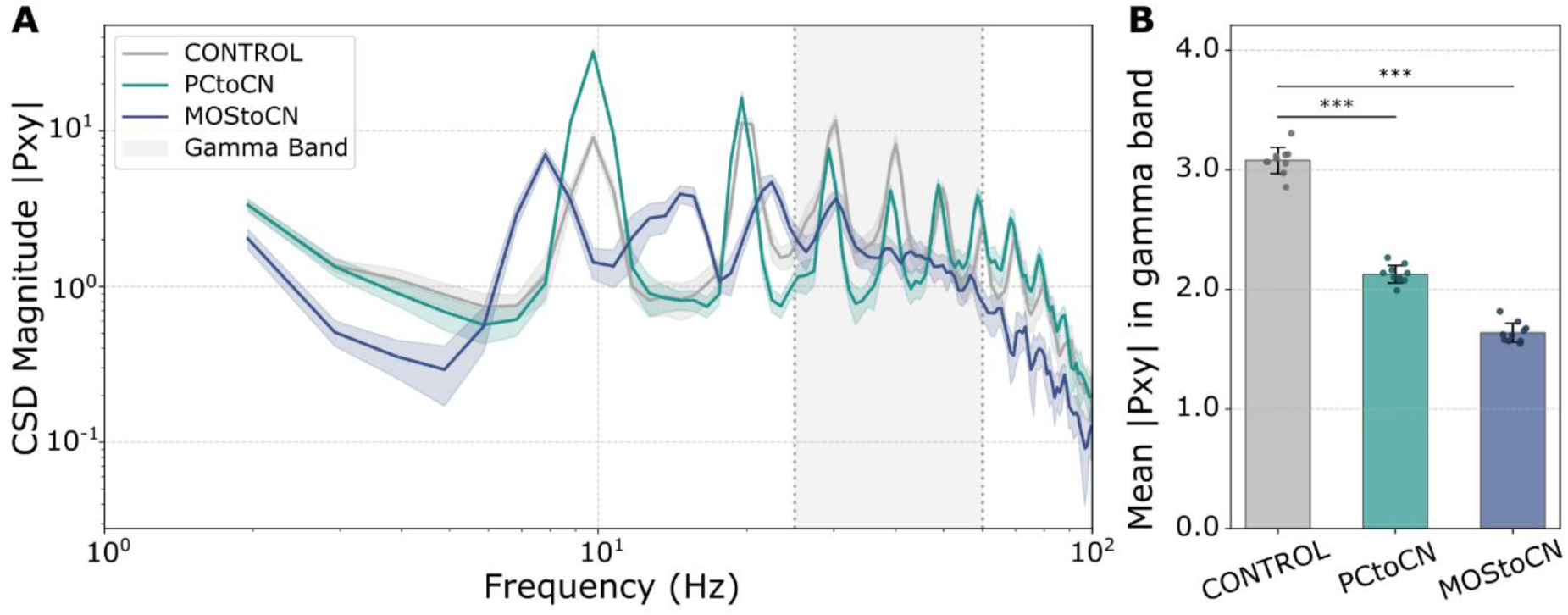
Cross-spectral density between mossy fibers and CNe population activity. **(A)** CSD magnitude across frequencies for CONTROL and CN input lesion conditions, represented as mean ± STD of 10 simulation runs for each condition, averaging over hemispheres. The gamma band is highlighted as the grey area. **(B)** Average CSD magnitude (|Pxy|) in the gamma band with comparisons between CONTROL and each lesion condition (***p < 0.001; independent-samples two-sided unpaired t-test, FDR-corrected over all lesion conditions). MOStoCN: inactivating the direct excitatory projection from mossy fibers to CNe; CNe: excitatory cerebellar nuclei neurons; CSD: cross-spectral density; PCtoCN: inactivating the indirect inhibitory projection from PC to CNe.

#### 2.3.4 Changes in cerebellar spiking activity with microcircuit lesions

Exploiting the higher granularity of the SNN model, we also analyzed mossy fibers, PC and CNe spiking activity in the cerebellar network, to identify microscale changes underlying the observed alterations of gamma-band power, coherence and CSD with the input to CNe lesions. To understand the spiking mechanisms underlying changes in population gamma power, we extracted measures of intra-population spiking activity, i.e., the phase-locking value (PLV) (Lowet et al., 2016) between the spikes of each CNe and the CNe population firing rate signal in the gamma band and the pairwise correlation of CNe spike trains as a measure of intra-population synchronization. The PLV of CNe was significantly decreased in both CN input lesions (Supplementary Fig. S8A), which is reflected in the changes in gamma-band power for the CNe population (Table 1): with these lesions, individual CNe spikes become less synchronized to the population gamma rhythm, decreasing the gamma-band power. Also, the overall CNe population synchronization decreased in both lesions (Supplementary Fig. S8B).

To explain the changes of oscillatory coupling between cerebellar input and output populations, as measured through coherence (Table 1) and CSD, we measured the PLV between CNe spikes and the mossy fiber firing rate signal filtered in the gamma band, showing that it was significantly reduced in *MOStoCN* and *PCtoCN* (Fig. 6A). Instead, cerebellar input-output population synchronization measured through cross-correlation of firing rate signals and pairwise cross-correlation of spike trains between mossy fibers and CNe, decreases with *MOStoCN* and increases with *PCtoCN* (Fig. 6B-C), explaining the observed changes in coherence with lesions to CN input (Fig. 4A-B) (Table 1), which go in the same direction.

**Figure 6.**
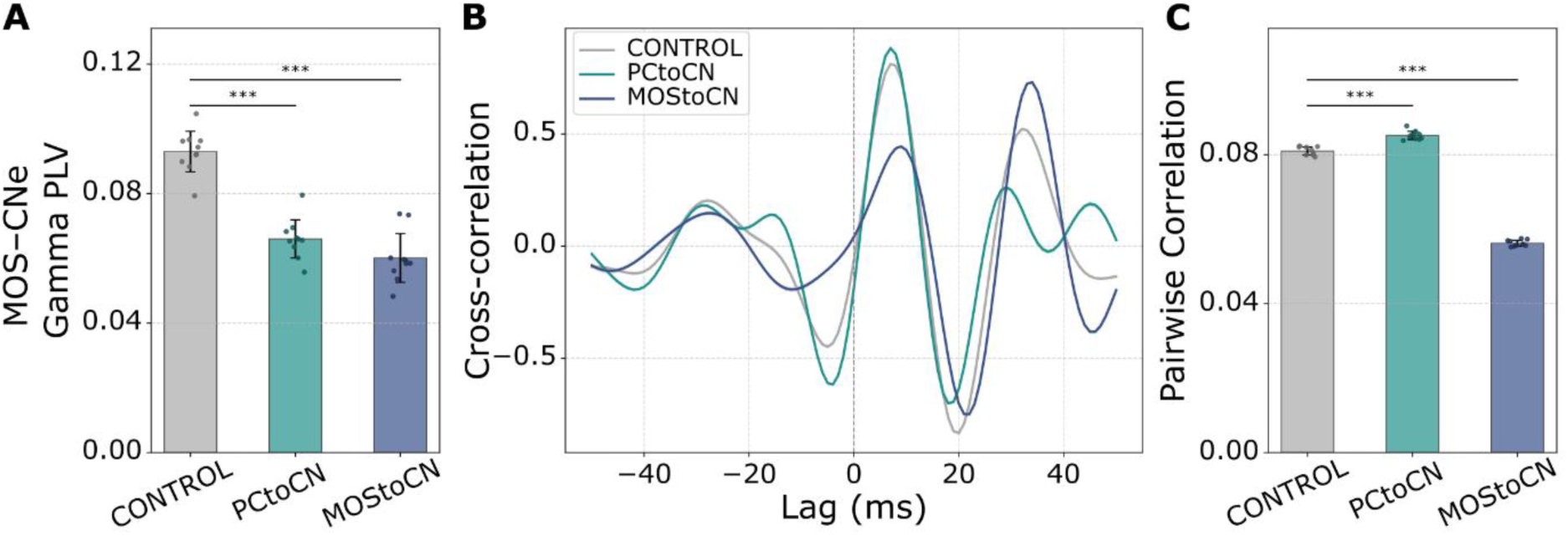
Cerebellar circuit spiking synchronization measures. **(A)** PLV of CNe spikes with respect to mossy fiber (MOS) firing rate in the gamma band, as mean ± STD across 10 simulation runs per condition. **(B)** Normalized cross-correlation between mossy fiber and CNe activity filtered in the gamma band, in CONTROL and lesion conditions, represented as mean ± STD of 10 simulation runs for each condition, averaging hemispheres. **(C)** Cerebellar spiking synchrony measured as pairwise spike train correlation between mossy fibers and CNe spikes in time bins of 20ms. Bar plots and error bars respectively show the mean and standard deviation over 10 simulations, each one averaged across the two hemispheres, with comparisons between CONTROL and each lesion condition (***p < 0.001; independent-samples two-sided unpaired t-test, FDR-corrected over all lesion conditions). PLV: Phase-locking value; CNe: excitatory cerebellar nuclei neurons; MOStoCN: inactivating the direct excitatory projection from mossy fibers to CNe; PCtoCN: inactivating the indirect inhibitory projection from PC to CNe.

## 3. Discussion

Our multiscale co-simulations provide a potential mechanism for how cerebellar microcircuitry drives macroscopic cortical dynamics during sensorimotor integration. Here, we show how the cerebellum may enhance M1-S1 gamma-band coherence during whisking not only thanks to the connections with the rest of the brain (Popa et al., 2013)(Fig. 2D-E), but also through the spike processing inside the cerebellar circuit, providing new insight into underlying mechanisms of sensorimotor integration. We found that synaptic inputs to the CN (from mossy fibers and PCs) differentially impact oscillations’ power and coherence by regulating the synchronization of spikes within the CN and sensorimotor cortex.

Modulation of the gamma-band by the cerebellum is akin to the fast dynamics of its circuit (D’Angelo, 2011). The cerebellum reacts to the inputs in about 10ms, i.e., one to two orders of magnitude faster than the cerebral cortex (Bower, 2002). Moreover, cerebellar control on movements operates in the millisecond range, reaching a precision unattainable in cerebro-cortical circuits (Timmann et al., 1999). Cerebellar gamma-band oscillations and cortical coherence have been hypothesized to be linked (De Zeeuw et al., 2008). In our model, gamma power and coherence seem functionally distinct (Table 1): a drop in local gamma power does not always mean that the network becomes less coherent; in cases like *PCtoCN*, the network becomes more synchronized despite lower gamma power.

Regarding power, we showed that the cerebellum acts as a ‘transformer’ of frequencies, enhancing gamma power and decreasing theta power along its circuit. In our model, gamma-band power matched experimental observations:

1. Both PC and CNe exhibit gamma-band oscillations (Fig. 3) (Middleton et al., 2008; Person & Raman, 2011).
2. Both mossy and PC inputs to CN are necessary to sustain gamma-band oscillations in the CN relative to the input (Fig. 3D), confirming that input signal processing in the cerebellar cortex, conveyed to the CN through PC, is essential for controlling the cerebellar output in the gamma band (Person & Raman, 2011).
3. MLI inhibition contributes to gamma-band oscillations in PC (Supplementary Fig. S6A) (Middleton et al., 2008).

Our model also offers novel insights into cerebellar processing. While PCs show increased gamma and theta power relative to mossy fibers, we observe an increase in gamma and decrease in theta power at the CNe (Fig. 3). When lesioning the connection between mossy fibers and CN, the gamma power increase is also significantly reduced in PCs, with small effect size though, probably because lesions in this cerebellar pathway involve other brain regions through recurrent loops reverberating on the cerebellum. Thus, both cerebellar pathways are important for the transforming power of the ‘cerebellar engine’.

Regarding coherence, our results point to different roles of cerebellar circuit elements not only in transferring but also in diversifying the cerebellar output signals to the cortex. The direct transmission of inputs from mossy fibers to CN appears necessary to keep cerebellar input-output and M1-S1 activity coherent at control levels (Fig. 4 and 6). In the indirect pathway through PCs, different mechanisms may be at play. Inactivation of the inhibitory connection from PC to CNe increased cerebellar and sensorimotor cortex coherence (Fig. 4) and synchronization between mossy fibers and CNe (Fig. 6). This result suggests that PC inhibition to CN might be more relevant for fine tuning the movement (e.g. velocity or acceleration adjustment) (S. Wu et al., 2024) than for overall sensorimotor integration, e.g., during baseline exploration or early adaptation. In these phases, the inhibitory PC input decorrelates CN firing patterns (Fig. 6) with respect to mossy fibers, thereby enhancing the information capacity and processing efficiency of the cerebellum. By decorrelating CN activity, Purkinje inhibition ensures that the CN output is not simply a redundant copy of the input, allowing for more efficient encoding of sensorimotor or cognitive information, essential for motor learning. On the other hand, in late adaptation conditions, the PC activity is reduced due to long-term depression plasticity mechanisms. Our model suggests that in this condition, when the CN receive less inhibitory input from PC, cerebellar input-output and (indirectly) M1-S1 coherence increase. Therefore, release of CN from PC inhibition during motor learning might be important not just for precise adaptation of movement kinematics (Heiney et al., 2014), but also for synchronization across brain regions, potentially improving the learning and binding processes that are essential to allow sensorimotor prediction. When analyzing gamma relative to theta coherence (Fig. 4), dual inputs to the CN are required to maintain balanced coherence across frequency bands, both in the cerebellar circuit and in the sensorimotor cortex. Further, mossy fiber excitation and Purkinje inhibition to the CN are important for phase-locking of cerebellar output spikes to the gamma-band component of the input signal, impacting gamma-band power and coherence combined, as observed through similar changes in CSD (Fig. 5).

When the PC input to CN is intact, MLI inhibition is necessary to keep cerebellar input and output coherent, even if changes in cerebellar coherence do not impact M1-S1 synchronization (Supplementary Fig. S6B-C). Thus, MLI inhibition is probably more important for cerebellar communication with other subcortical structures than for sensorimotor cortical control (Gaffield & Christie, 2017). It could also play a pivotal role during reflexive whisking (Brown et al., 2025) rather than free whisking and may be acting on frequency bands other than those investigated here.

Reproducing these complex, task-driven macroscopic dynamics required overcoming significant modeling challenges. Unlike typical resting-state models, our framework successfully sustained multi-frequency oscillations during an active behavioral paradigm (whisking), by integrating a task-specific pathway gain, with state-of-the art methods, like thalamocortical loop neural masses and indegree-based homeostatic feedback inhibition (Stasinski et al., 2024). While previous models have utilized the cerebellum as an isolated module for neurorobotic control, we embedded a realistic spiking neural network (SNN) within a whole-brain architecture, capturing the complex, recurrent loops between the cerebellum and the sensorimotor cortices. For example, results demonstrate that cutting a connection in this loop does not necessarily bring about a decrease in power, or lesioning different pathways alters the gamma-to-theta power ratio in specific ways. The recurrent loops between the mean-field model and the SNN cause lesions in the SNN to impact the rest of the brain and then, indirectly, modulate the SNN itself through changes in mossy fiber oscillatory activity. This phenomenon explains, e.g., why *MOStoCN* affects PC properties (e.g. Fig. 3C). In addition, the dynamical properties of neural population activity in the SNN are emerging properties, not constrained during fitting (<u>Par. 2.2</u>), but resulting from the data-driven reconstruction (De Schepper et al., 2022). Finally, due to the intrinsic closed-loop nature of the model, fitting was not straightforward as in single-scale models since parameter choices on different scales are highly interdependent. Here, we introduced a first comprehensive reusable pipeline to fit model parameters at the different scales against multimodal experimental datasets, using simulation-based inference (Tejero-Cantero et al., 2020). A first follow-up study demonstrates the versatility of our framework by investigating the use case of the spread of beta oscillations in Parkinson’s disease (Gambosi et al., 2026). Taken together, the developed and validated comprehensive model bridges the scales in a biologically realistic way, opening avenues of exploring various multiscale animal and future human behavioral paradigms.

While our model captures critical sensorimotor dynamics, future refinements will expand its applicability. Currently, the structural connectome relies on cerebello-thalamo-cortical loops mapped to sensorimotor areas according to the AMBCA. However, particularly in humans, a vast majority of cerebellar projections target associative regions like the prefrontal cortex nuclei (Pisano et al., 2021). Updating the whole-brain connectome to reflect these pathways will allow the model to investigate the cerebellum’s impact on cortical dynamics during higher cognitive tasks. Furthermore, our current whisking paradigm relies on a minimal pathway gain. Incorporating parallel sensorimotor loops in future iterations will be critical to fully resolve how distributed network architectures influence cerebello-cortical synchronization. While we focused on free whisking, the framework could be extended to reflexive whisking, where climbing fibers (the other input to the cerebellar circuit) also play a crucial role (Romano et al., 2018).

Ultimately, this multiscale framework provides a mechanistic tool for investigating neurological and neurodevelopmental disorders. Pathologies such as cerebellar stroke or autism spectrum disorder are frequently characterized by localized microcircuit alterations, such as disrupted PC excitability or synaptic transmission. Yet, clinical symptoms often emerge as widespread large-scale functional dysconnectivity (Riva et al., 2013). By bridging cellular perturbations to system-level dynamics, our framework enables the virtual lesioning of specific microcircuits, offering a unified platform to reconcile cellular anomalies with the macroscopic functional deficits observed in patient populations.

## 4. Methods

### 4.1 Whole-brain connectivity

#### 4.1.1 Generating the connectome

The TVMB connectome was based on the AMBCA (Oh et al., 2014) and extracted following (Melozzi et al., 2017). For each hemisphere, 298 areas (Supplementary Table S1) were included, with the ratio between projection and injection density as connection strength, in a grid volume of 100-μm-side voxels (Oh et al., 2014). The left-hemispheric connectome was built mirroring the right-hemispheric one, given the high lateral symmetry in mouse brain connections (Calabrese et al., 2015), resulting in a connectome with 596 regions. We set self-connections to zero. Tract lengths were calculated as the Euclidean distance between the centers of each region pair, taken from the Allen Software Development kit (https://allensdk.readthedocs.io/en/latest/), following (Melozzi et al., 2017).

#### 4.1.2 Merging regions into major structures and distributing specific thalami

Regions in the connectome were assigned to one of twelve major structures following the Allen reference ontology (*thalamus*, *cortical subplate*, *hippocampal formation*, *hypothalamus*, *medulla*, *midbrain*, *pallidum*, *striatum*, *olfactory areas*, *cerebellum*, *isocortex*, *pons* (connectivity.brain-map.org; (Dong, 2008; Oh et al., 2014), Supplementary Table S1). Regions not expected to be involved in whisking pathways (Bosman et al., 2011) were merged into their respective major structure in each hemisphere, thus reducing the complexity and size of the whole-brain network, without losing explanatory power for our specific research question.

For the *thalamus*, we distinguished between specific and non-specific thalamic nuclei (Van Der Werf et al., 2002; White, 1979). For the specific ones, we merged the *ventral anterior-lateral nucleus* and the *ventral posteromedial nucleus* regions. Then we substituted the merged regions with 43 specific thalamic nuclei nodes per hemisphere with the same center point coordinates, each one connected to one isocortical region unilaterally, forming thalamocortical loops, as well as to a corresponding reticular thalamic part. For the non-specific thalamus, the *midline* and *intralaminar thalamic nuclei* were merged (Van Der Werf et al., 2002) and connected to all regions (except for specific thalamic ones) based on the connectome.

For merged regions, the connectivity weights were summed up, while tract lengths were computed as averages of the original tract lengths, weighted by the respective connection weight. As a result, we obtained a whole-brain connectome with 214 regions (Supplementary Table S2): (i) the *isocortex*, i.e., 43 unmerged isocortical regions per hemisphere, each one with its own specific thalamic nucleus; (ii) the non-isocortex for the whisking subnetwork (Fig. 1A-B), i.e., selected regions from *pons* (*pons motor*, *pons behavioral*, PrV from the *pons sensory*, and the rest of the *pons sensory*), *medulla* and *midbrain* [IO, *spinal nucleus of the trigeminus (sensory afferences)* (SpV), *facial motor nucleus (motor efferences)* (FMN), and the *superior colliculus (motor related)*, and the rest merged]; (iii) the *cerebellum* with the AN selected from *cerebellar cortex* and the rest merged, and all CN (dentate, interposed, fastigial) merged; (iv) the major non-isocortical structures outside of the whisking circuit merged, i.e. *cortical subplate*, *hippocampal formation*, *hypothalamus, pallidum, striatum, and olfactory areas*.

#### 4.1.3 Connectivity normalization and delays’ lower bound

The resulting connectivity weights were log-transformed and normalized (Triebkorn et al., 2024) following

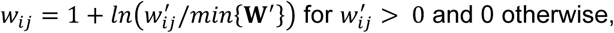

where *w*^′^_*ij*_ and *w_ij_* are the weights before and after normalization, respectively, and *min*{**W**^′^} denotes the minimum non-zero value of the connectivity matrix before normalization. We further divided the resulting weights by their 99^th^ percentile. Transmission speed along connections was set to 3m/s and a lower bound was set for tract lengths, such that the minimum delay was equal to one TVMB integration time step (0.1ms), resulting in transmission delays *τ_ij_* for all node pairs.

After normalization, all weights between isocortical and their respective specific thalamic nodes were set to 1. All connections between specific thalamic nodes and all other nodes were set to 0, except for whisking-specific connections, i.e., the contralateral connections from SpV to the specific thalamic nucleus of S1 and from CN to the specific thalamic nuclei of M1 and of S1 (Bosman et al., 2010, 2011). Subsequently, two whisker nodes (left and right “whiskers”) were added to close the sensorimotor loop between the brain and the neural periphery. All connectivity for these two nodes was set to 0 except for the ipsilateral incoming connections from FMN (conveying the motor command from M1) and outgoing connections to SpV (for proprioception), which were both set to 1. The respective time delays were set to 1ms.

#### 4.1.5 Whisking-specific pathway

For the simulated whisking activity to be transmitted via the sensorimotor loop passing through the cerebellum, connectivity weights along a minimal whisking-specific pathway were strengthened (Fig. 1B) based on the whisking-specific subnetwork of (Bosman et al., 2010): (a) from M1 to FMN contralaterally and (b) from FMN to “whisker” ipsilaterally for the motor command (Sreenivasan et al., 2016), (c) from “whisker” to SpV ipsilaterally for proprioception, (d) from SpV to PrV, (e) from PrV to AN and (f) from AN to CN ipsilaterally, for the path via cerebellum, and, finally, (g) from the CN to the specific thalamic nuclei of M1 and (h) to the specific thalamic nucleus of S1, contralaterally, for the sensorimotor loop to close. The chosen connections were strengthened following *g_ij_w_ij_*, where *w_ij_* is the connection weight between source region *j* and target region *i* and *g_ij_* ∈ [1, 99] is the gain, which was constrained such that *w*^′^ ⁄*D_i_* ≤ 0.99, where *D_i_* = ∑*_j_ w_ij_* is the indegree of target region *i*. In all cases, except for the last two connections (g) and (h), all weights of the remaining connections towards the target region *i* were reduced to maintain the same indegree *D_i_* (Supplementary Note S1.1 and Supplementary Fig. S8 for details on how *g_ij_* were optimized against M1-S1 coherence drop data from (Popa et al., 2013) with *cerebOFF* and Supplementary Table S5 for the resulting fitted parameter values).

### 4.2 The multiscale virtual mouse brain

#### 4.2.1 TVMB model dynamics

The typical summary equation of a stochastic TVB model (Ritter et al., 2013; Sanz-Leon et al., 2015) including additive noise is

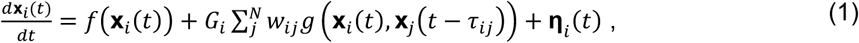

where **x***_i_*(*t*) denotes the vector of state variables describing the dynamics of a region node *i*, *f*(.) is the function of the node dynamics, *g*(.) is the coupling function between any pair of nodes *i* and *j*, *w_ij_* is the coupling strength, *G_i_* is a parameter scaling the coupling for each node *i* and, finally, **η***_i_*(*t*) is an optional vector of stochastic inputs, typically white noise **η***_i_*(*t*)∼*Normal*(0, **σ*_i_***), i.e., samples of a normal distribution with mean 0 and standard deviation **σ*_i_***.

The neural mass model used for TVMB nodes was a modification of the cortico-thalamic Wilson-Cowan model from (Griffiths et al., 2020; Griffiths & Lefebvre, 2019), which was based on previous models of (Breakspear et al., 2006; Freyer et al., 2011; Robinson et al., 2001). This model allows to reproduce biologically plausible thalamocortical loop dynamics of rodent sensorimotor cortices (Marshall et al., 2016), exhibiting oscillations in the gamma band modulated by oscillations in lower-frequency bands (e.g. theta for rodent models (Griffiths et al., 2020)).

Thalamocortical loops include isocortical nodes with excitatory and inhibitory populations (Fig. 1C) governed by

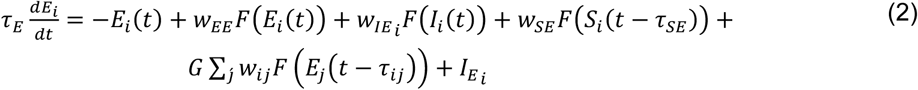

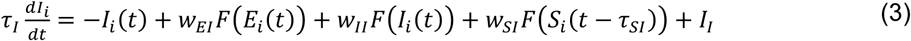

and specific thalamic nodes with relay and reticular population with differential equations

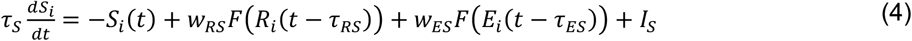

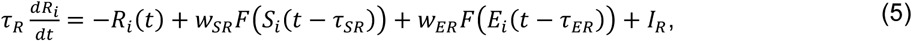

where *E_i_*, *I_i_*, *S_i_* and *R_i_* represent the state variables of the excitatory isocortical, inhibitory isocortical, excitatory relay and inhibitory reticular specific thalamic populations, respectively, whereas the index *i* denotes the isocortical region and the respective connected specific thalamic nucleus. The index *j* denotes all other regions that are not specific thalamic regions (denoted as “(non-)isocortical” from now on) that form long-range connections with an isocortical region *i*.

Non-isocortical nodes include one excitatory and one inhibitory neural population (Fig. 1C) with dynamics

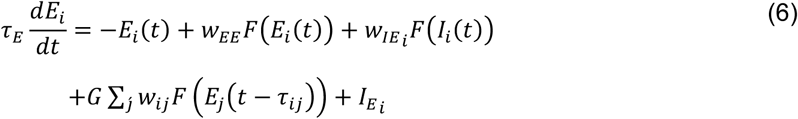

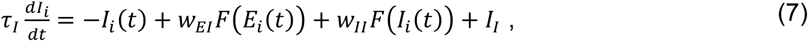

where *E_i_* and *I_i_* represent the state variables of the excitatory and inhibitory non-isocortical populations, respectively, and the index *j* denotes all other (non-)isocortical regions that form long-range connections with a non-isocortical region *i* (including the ipsilateral “whisker” node).

Finally, whisker nodes include one excitatory neural population per hemisphere *W_i_* with dynamics

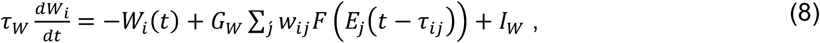

where *W_i_* receives only one long-range connection from the ipsilateral FMN conveying the motor command.

In Equations (2-8), 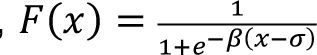 is a sigmoidal activation function, *I_E_*, *I_I_*, *I_S_*, *I_R_* and *I_W_* are constant baseline current parameters, *τ_*E*_*, *τ_I_*, *τ_S_*, *τ_R_* and *τ_W_* are time scale constants, *G* and *G_W_* (specific for whisker nodes) denote long-range coupling scaling parameters, and, finally, all other *w_XY_* and *τ_XY_* parameters denote connectivity weights and delays, respectively, between populations *X* and *Y*, where *X* and *Y* take values following the state variables (*E*, *I*, *S* and *R*). In that respect, TVMB long-range connectivity weights and delays between each isocortical region and its respective specific thalamic nuclei were overwritten with the respective *w_XY_* and *τ_XY_* of the original thalamocortical neural mass model of (Griffiths et al., 2020; Griffiths & Lefebvre, 2019). Parameter *I_S_* was optimized against empirical data of M1 and S1 PSDs from (Popa et al., 2013) (Supplementary Note S1.1 and Supplementary Fig. S9 for details and Supplementary Table S4 for the resulting fitted parameters’ values), whereas parameters *I_W_* and *G_W_* against M1-S1 coherence drop data from (Popa et al., 2013) with inactivated cerebellum, together with parameters *g_ij_* (<u>Par. 4.1.5</u>; Supplementary Note S1.1 and Supplementary Fig. S10 for details and Supplementary Table S5 for the resulting fitted parameter values).

#### 4.2.2 Scaling excitation – inhibition balance with indegree connectivity

To prevent over-excitation and achieve biologically plausible regional activities for a wide range of values of the global coupling *G*, we implemented a mechanism inspired by feedback inhibition control (Deco et al., 2014;) based on modulating the inhibitory weight from the inhibitory population to the excitatory one. We customized the baseline current parameter of excitatory populations (*I_E_*) as well as the local inhibitory weight (*w_IE_*) of each region, by scaling them based on an individual indegree scaler (Ódor et al., 2022; Rocha et al., 2018)

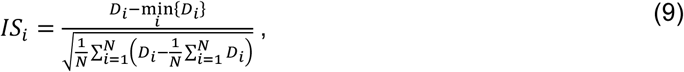

where *N* is the number of all isocortical and non-isocortical regions (i.e., excluding specific thalamic regions) and min{*D_i_*} the minimum value across all *D_i_*. The final parameter values for each region *i* are given by the equations

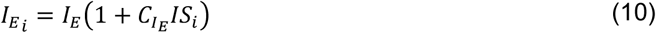

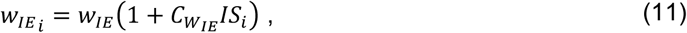

where *I_E_* = −0.35 and *w_IE_* = −3.0 as in the original model, *C_IE_* and *C_WIE_* are additional scaling parameters, which we fit to M1 and S1 PSDs from (Popa et al., 2013) together with parameter *I_S_* (Supplementary Note S1.1 and Supplementary Fig. S9).

### 4.3 Spiking neural network model

For multiscale co-simulations, the olivocerebellar circuit within the whole-brain network was modelled and simulated as an SNN in NEST3 (Hahne et al., 2021) (Fig. 1D), including (i) a detailed cerebellar cortex as the AN (Chen et al., 2016), reconstructed based on geometrical and morphological features of local microcircuitry (De Schepper et al., 2022) and (ii) a corresponding model of CN and IO circuits, to form an olivocerebellar microcomplex (Geminiani et al., 2024). In each layer, the main neural populations were included, namely granule cells and Golgi cells receiving inputs from mossy fibers, MLIs including basket cells and stellate cells, PCs, CNe and inhibitory CN neurons, IO neurons (Fig. 1D). Single neurons were modelled as extended-generalized leaky integrate and fire capturing the complex non-linear dynamics of cerebellar neurons, with parameters optimized to fit population-specific electro-responsive properties (Geminiani, Casellato, et al., 2019; Geminiani et al., 2018). Synapses were modeled as alpha-shaped conductance-based, with weights set to reproduce baseline *in vivo* activity (Rancz et al., 2007; ten Brinke et al., 2015, 2017) and time constants and connection delays from literature (Cavallari et al., 2014; Geminiani et al., 2024; Geminiani, Pedrocchi, et al., 2019). Further, neurons in the PrV (from *pons sensory*) were modelled as spiking units, relaying spike trains from the whisking input regions to the corresponding mossy fibers also modelled as relay units in the olivocerebellar SNN.

### 4.4 Inter-scale interface models

Co-simulation of the multiscale model requires interfaces for the transformation and exchange of activity between the TVMB neural mass nodes and the NEST SNN. For the TVMB to NEST interface, the TVMB’s total large-scale coupling activity *C_i_*(*t*) was converted to a rate via a linear transformation *r_i_^TVMB^*^→*NEST*^(*t*) = *w^TVMB^*^→*NEST*^*C_i_*(*t*), where *C_i_*(*t*) = *G* ∑*_j_ w_ij_ F* (*E_j_*(*t* − *τ_ij_*)), *i* is the index of the SNN target regions (neurons in the PrV and mossy fibers in the AN of each hemisphere) and *j* the index of all (non-)isocortical regions coupling with each one of them, whereas *w^TVMB^*^→*NEST*^ is a weight to be tuned aiming for a firing rate of 4.2Hz for the granule cells of AN (Chen et al., 2017) (Supplementary Note S1.2 and Supplementary Fig. S2). This rate was converted to a different spike train for each neuron in the target population of the SNN input regions (Fig. 1D), according to a Poisson process, and delivered with a delay equal to the minimum NEST simulation delay (i.e., 0.05ms). For the NEST to TVMB interface, the instantaneous average firing *r^NEST^*^→*TVB*^(*t*) was computed from population spikes for the output population of each brain region *i* modelled in NEST (Fig. 1D). Then, the TVMB activity *E*_i_ (*t*) was updated via linear transformation and integration *τ* 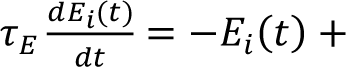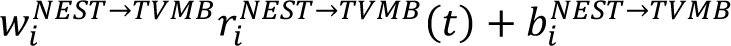, acting as a low-pass filter of the spiking activity, using the same integration scheme and time step as for the TVMB simulation (<u>Par. 4.5</u>). Parameters *b^NEST^*^→*TVMB*^ and *w^NEST^*^→*TVMB*^ of the linear transformation were also tuned for the output TVMB signal to match the respective signals’ baselines and amplitudes of TVMB-only simulations.

### 4.5 Simulations

We performed two kinds of simulations: (a) of the large-scale neural mass TVMB model alone (“TVMB-only”), in which case all cerebellar subcortical regions were modelled identically to the rest of the subcortical regions (<u>Par. 4.2.1</u>) and (b) co-simulations (“TVMB-NEST”) of the multiscale TVMB-NEST model that included the SNN olivocerebellar circuit. To validate the model against the experimental observations of (Popa et al., 2013), we simulated 10 runs under four conditions: (i) TVMB-only simulations with cerebON, (ii) TVMB-only simulations with cerebOFF, (iii) TVMB-NEST co-simulations with cerebON, and (iv) TVMB-NEST co-simulations with cerebOFF.

For all such (co-)simulations, a stochastic Euler-Maryuama integration method with a time step of *dt* = 0.1ms and additive white noise of standard deviation equal to σ*_i_* = 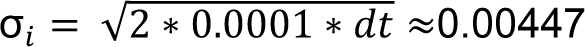 ≈0.00447 was used for the TVMB component of the simulation. Initial conditions for the neural mass TVMB model state variables up to one second in the past were chosen from a normal distribution with mean and standard deviation equal to the baseline currents (*I*_{_*_E_*_,*I*,*S*,*R*}_) of each population *i*, i.e., *SV_i_*(*t*_0_)∼*Normal*(*I_SV_*, *I_SV_*), for all times *t*_0_ ∈ [−1000, 0]ms and state variables *SV* ∈ {*E*, *I*, *S*, *R*} (specifically for the excitatory population *E*, the mean value of *I_E_* over all regions was used), except for the whisker nodes, for which *W_i_*(*t*_0_) = 0. All source TVMB signals were recorded with a sampling period of 1ms.

For the NEST component of simulations, the integration time step and minimum delay of NEST was set to half the TVMB time step, i.e., 0.05ms. The membrane potential of each neuron was initialized with *E*_L_ + *Uniform* 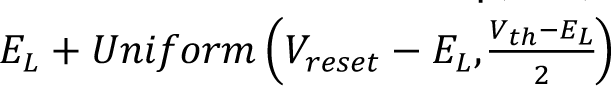, where *E*, *V* and *V* are leakage, reset and threshold potential, respectively. For cerebOFF simulations, all connections to and from AN, the rest of cerebellar cortex, and CN were set to 0 and in co-simulations the threshold potential of the CNe neurons (output population in the cerebellum) was raised to *V_t_*_ℎ_ = −35.0*mV*.

### 4.6 Microcircuit lesion co-simulations

We applied localized lesions within the SNN in TVMB-NEST co-simulations and quantified the changes in M1-S1 coherence. Specifically, we selectively inactivated synaptic connections within the cerebellar SNN: (a) mossy fibers projections to CNe, i.e. the direct nuclear pathway (*MOStoCN*); (b) inhibitory PC projections to CNe, i.e. the indirect cortical pathway (*PCtoCN*); (c) inhibitory projections from MLIs to PCs (*MLItoPC*), and (d) self-inhibition among MLIs (*MLItoMLI*), by setting the respective SNN connection weight to 0. To isolate synaptic effects on spiking dynamics without altering average firing rates (Heinzle et al., 2007), we adjusted the target population endogenous current parameter, which controls spontaneous firing (Geminiani et al., 2018). The new value was set to preserve the average firing rate of the target population and the CNe in 1s resting-state NEST-only simulations of the SNN, taken as the reference value (Supplementary Table S8).

To further control for activity baseline and amplitude confounds in whole-brain whisking co-simulations, we performed co-simulations also with normalized cerebellar output. Specifically, the NEST to TVMB interface parameters were adjusted such that the output CN to TVMB signal was normalized by its median baseline (50^th^ percentile) and amplitude (99^th^–50^th^ percentile) as measured in the corresponding *CONTROL* co-simulations. In all lesion conditions, cerebellar input-output and M1-S1 coherence were compared to values in *CONTROL* co-simulations.

### 4.7 Analysis and Statistics

From the excitatory activity signal of M1 and S1 nodes, we computed the PSD as the spectrogram normalized by the power across all frequencies and the spectral coherence between them. To compare cerebON and cerebOFF, we extracted the average value in the gamma band (25 – 60Hz) in both conditions and its drop as percentage of the normalized difference (*cerebON* – *cerebOFF*) / *cerebON*.

In the SNN virtual lesion co-simulations, for all conditions, we also computed coherence between cerebellar in- and output, from the activity of TVMB interface nodes of the trigeminal nucleus, conveying the whisking input signal to cerebellar mossy fibers and the CN, sending the output of the cerebellum to the rest of the brain. For both pairs of regions, we report the gamma-band and gamma-to-theta average values.

Regarding spiking activity, we computed the PSD of mossy fibers, PC and CNe population activity. We quantified the power of PC/CNe neurons relative to mossy fibers as power ratio, defined as the ratio between the integrals of the PSD for each of the two populations of interest (PC or CNe) and the mossy fibers in specific frequency bands.

Finally, as a measure of inter-population synchronization of spiking activity in the gamma band, we computed the PLV between CNe spikes and the mossy fiber population activity signal filtered in the gamma band, using the Hilbert transform to compute the instantaneous phase, the cross-correlation between mossy fibers and CNe firing rate signals, and the pair-wise correlation of spike trains in bins of 10ms. For internal population synchronization, we computed the PLV between CNe spikes and the overall CNe population activity signal filtered in the gamma band and the pairwise spike train correlations of CNe and PC populations in time bins of 10ms.

For all conditions and all extracted values, we report mean and STD over 10 simulation runs after averaging values for the two hemispheres in each run. For comparison between conditions, independent-samples two-sided unpaired t-test is used, with *p* < 0.05, *p* < 0.01, and *p* < 0.001 for statistical significance, applying FDR correction in the virtual lesion conditions, as multiple comparisons are run.

### 4.8 Hardware and software specifications

Simulations and fitting using simulation-based inference (SBI) have been performed on the High-Performance Cluster for Research and Clinic of the Berlin Institute of Health, Berlin, Germany.

### 4.9 Data and code availability

All code is available at https://github.com/virtual-twin/tvb-multiscale-rising-net/releases/tag/Geminiani_Meier_Perdikis_etal and can be run in the Docker image https://hub.docker.com/repository/docker/dionperd/tvb-multiscale-dev/tags/rising-net_publication/sha256:511c1be7209e0966948a588ca4f450932054e985dc17af96dc5830fa3c9c04aa. All data for reproducing results are uploaded in the repository https://zenodo.org/records/20530965.

## Acknowledgements

We thank Fulvia Palesi for initial discussions and Michael Schirner for insightful discussions on feedback inhibition control.

The authors used Claude by Anthropic for code scaffolding of the simulation data analysis and Google’s Gemini for code scaffolding for plotting. All AI-generated outputs were rigorously reviewed, edited and validated by the authors, who maintain full accountability for the accuracy and integrity of the final content.

PR acknowledges support by the EU Horizon Europe program: BRIDGE (101219311), EBRAINS2.0 (101147319), Virtual Brain Twin (101137289), EBRAINS-PREP 101079717, AISN 101057655, EBRAIN-Health 101058516, EIC grant PHRASE 101058240, the Digital Europe Programme: TEF-Health (101100700), SHAIPED (101195135), CoordinaTEF (101168074), the German Research

Foundation: SFB 1436 (project ID 425899996); SFB 1315 (project ID 327654276); SFB 936 (project ID 178316478; SPP Computational Connectomics RI 2073/6-1, RI 2073/10-2, RI 2073/9-1; DFG Clinical Research Group BECAUSE-Y 504745852, DFG German-Spanish project BrainsAge RI 2073/14-1, DFG Project-ID 424778381 - TRR 295 Retune, ERA PerMed JTC2021 German Federal Ministry for Health PatternCog ZMI5-2522FSB904, Berlin University Alliance OpenMake, the Virtual Research Environment at the Charité Berlin and EBRAINS Health Data Cloud and the Berlin Institute of Health and Foundation Charité. JM acknowledges support by the Deutsche Forschungsgemeinschaft (DFG,

German Research Foundation) - Project-ID 424778381 - TRR 295. This research has received a Voucher (CEoI 4-Rodent microcircuits: RisingNet Whole-bRaIn rodent SpikING neural NETworks) from the European Union’s Horizon 2020 Framework Programme for Research and Innovation under the Specific Grant Agreement No. 945539 (Human Brain Project SGA3) to EDA, supporting AG, CC and ED. AG also acknowledges support by IBRO through an IBRO Short Stay Grant. ED acknowledges support by the EU Horizon Europe program: EBRAINS2.0 (101147319), Virtual Brain Twin (101137289), TEF-Health (101100700), and by the NEXTGENERATIONEU (NGEU) and funded by the Ministry of University and Research (MUR), National Recovery and Resilience Plan (NRRP), project MNESYS (PE0000006) – A Multiscale integrated approach to the study of the nervous system in health and disease (DN. 1553 11.10.2022), and project EBRAINS-Italy (PIR0000011, CUP B51E22000150006, “EBRAINS-Italy”). CC acknowledges support by EU Horizon Europe program EBRAINS2.0 (101147319).

## References

1. Abbruzzese, G., & Berardelli, A. (2003). Sensorimotor integration in movement disorders. Movement Disorders, 18(3), 231–240. 10.1002/MDS.10327

2. Bosman, L. W. J., Houweling, A. R., Owens, C. B., Tanke, N., Shevchouk, O. T., Rahmati, N., Teunissen, W. H. T., Ju, C., Gong, W., Koekkoek, S. K. E., & De Zeeuw, C. I. (2011). Anatomical pathways involved in generating and sensing rhythmic whisker movements. Frontiers in Integrative Neuroscience, 5, 53. 10.3389/fnint.2011.00053

3. Bosman, L. W. J., Koekkoek, S. K. E., Shapiro, J., Rijken, B. F. M., Zandstra, F., van der Ende, B., Owens, C. B., Potters, J.-W., de Gruijl, J. R., Ruigrok, T. J. H., & De Zeeuw, C. I. (2010). Encoding of whisker input by cerebellar Purkinje cells. The Journal of Physiology, 588(Pt 19), 3757–3783. 10.1113/jphysiol.2010.195180

4. Bower, J. M. (2002). The organization of cerebellar cortical circuitry revisited: implications for function. Annals of the New York Academy of Sciences, 978, 135–155. 10.1111/J.1749-6632.2002.TB07562.X

5. Breakspear, M., Roberts, J. A., Terry, J. R., Rodrigues, S., Mahant, N., & Robinson, P. A. (2006). A unifying explanation of primary generalized seizures through nonlinear brain modeling and bifurcation analysis. Cerebral Cortex, 16(9), 1296–1313. 10.1093/cercor/bhj072

6. Brown, S. T., Holla, M. R., Land, M. A., Yang, S., McDonald, A. J., St-Pierre, F., & Raman, I. M. (2025). High-speed voltage imaging of action potentials in molecular layer interneurons reveals sensory-driven synchrony that augments movement. Cell Reports, 44(8), 116148. 10.1016/j.celrep.2025.116148

7. Calabrese, E., Badea, A., Cofer, G., Qi, Y., & Johnson, G. A. (2015). A Diffusion MRI Tractography Connectome of the Mouse Brain and Comparison with Neuronal Tracer Data. Cerebral Cortex, 25(11), 4628–4637. 10.1093/cercor/bhv121

8. Cavallari, S., Panzeri, S., & Mazzoni, A. (2014). Comparison of the dynamics of neural interactions between current-based and conductance-based integrate-and-fire recurrent networks. Frontiers in Neural Circuits, 8(March), 1–23. 10.3389/fncir.2014.00012

9. Chen, S., Augustine, G. J., & Chadderton, P. (2016). The cerebellum linearly encodes whisker position during voluntary movement. ELife, 5, e10509. 10.7554/eLife.10509

10. Chen, S., Augustine, G. J., & Chadderton, P. (2017). Serial processing of kinematic signals by cerebellar circuitry during voluntary whisking. Nature Communications, 8(1), 232. 10.1038/s41467-017-00312-1

11. D’Angelo, E. (2011). Neural circuits of the cerebellum: hypothesis for function. Journal of Integrative Neuroscience, 10(3), 317–352. 10.1142/S0219635211002762

12. De Schepper, R., Geminiani, A., Masoli, S., Rizza, M. F., Antonietti, A., Casellato, C., & D’Angelo, E. (2022). Model simulations unveil the structure-function-dynamics relationship of the cerebellar cortical microcircuit. Communications Biology 2022 5:1, 5(1), 1240-. 10.1038/s42003-022-04213-y

13. De Zeeuw, C. I., Hoebeek, F. E., & Schonewille, M. (2008). Causes and Consequences of Oscillations in the Cerebellar Cortex. Neuron, 58(5), 655–658. 10.1016/j.neuron.2008.05.019

14. Deco, G., Ponce-Alvarez, A., Hagmann, P., Romani, G. L., Mantini, D., & Corbetta, M. (2014). How local excitation-inhibition ratio impacts the whole brain dynamics. The Journal of Neuroscience, 34(23), 7886–7898. 10.1523/JNEUROSCI.5068-13.2014

15. Dong, H. W. (2008). The Allen reference atlas: A digital color brain atlas of the C57Bl/6J male mouse. John Wiley & Sons Inc. https://psycnet.apa.org/record/2008-01792-000

16. Flanders, M. (2011). What is the biological basis of sensorimotor integration? Biological Cybernetics, 104(1–2), 1. 10.1007/S00422-011-0419-9

17. Freyer, F., Roberts, J. A., Becker, R., Robinson, P. A., Ritter, P., & Breakspear, M. (2011). Biophysical mechanisms of multistability in resting-state cortical rhythms. The Journal of Neuroscience, 31(17), 6353–6361. 10.1523/JNEUROSCI.6693-10.2011

18. Gaffield, M. A., & Christie, J. M. (2017). Movement Rate Is Encoded and Influenced by Widespread, Coherent Activity of Cerebellar Molecular Layer Interneurons. The Journal of Neuroscience, 37(18), 4751. 10.1523/JNEUROSCI.0534-17.2017

19. Gambosi, B., Perdikis, D., Meier, J., Geminiani, A., Antonietti, A., Mazzoni, A., Ferrigno, G., Ritter, P., & Pedrocchi, A. (2026). Dopamine Depletion Drives Whole-Brain Oscillatory Disruptions via Cortico–Subcortical Resonance: A Multiscale Model of Parkinson’s Disease in Mice. BioRxiv, 2026.05.15.725133. 10.64898/2026.05.15.725133

20. Geminiani, A., Casellato, C., Boele, H. J., Pedrocchi, A., De Zeeuw, C. I., & D’Angelo, E. (2024). Mesoscale simulations predict the role of synergistic cerebellar plasticity during classical eyeblink conditioning. PLOS Computational Biology, 20(4), e1011277. 10.1371/JOURNAL.PCBI.1011277

21. Geminiani, A., Casellato, C., D’Angelo, E., & Pedrocchi, A. (2019). Complex electroresponsive dynamics in olivocerebellar neurons represented with extended-generalized leaky integrate and fire models. Frontiers in Computational Neuroscience, 13. 10.3389/fncom.2019.00035

22. Geminiani, A., Casellato, C., Locatelli, F., Prestori, F., Pedrocchi, A., & D’Angelo, E. (2018). Complex dynamics in simplified neuronal models: reproducing Golgi cell electroresponsiveness. Frontiers in Neuroinformatics, 12(88), 1–19. 10.3389/fninf.2018.00088

23. Geminiani, A., Pedrocchi, A., D’Angelo, E., & Casellato, C. (2019). Response Dynamics in an Olivocerebellar Spiking Neural Network With Non-linear Neuron Properties. Frontiers in Computational Neuroscience, 13. 10.3389/fncom.2019.00068

24. Griffiths, J. D., & Lefebvre, J. R. (2019). Shaping Brain Rhythms: Dynamic and Control-Theoretic Perspectives on Periodic Brain Stimulation for Treatment of Neurological Disorders. In V. Cutsuridis (Ed.), Multiscale models of brain disorders (Vol. 13, pp. 193–205). Springer International Publishing. 10.1007/978-3-030-18830-6_18

25. Griffiths, J. D., McIntosh, A. R., & Lefebvre, J. (2020). A Connectome-Based, Corticothalamic Model of State- and Stimulation-Dependent Modulation of Rhythmic Neural Activity and Connectivity. Frontiers in Computational Neuroscience, 14, 575143. 10.3389/fncom.2020.575143

26. Hahne, J., Diaz, S., Patronis, A., Schenck, W., Peyser, A., Graber, S., Spreizer, S., Vennemo, S. B., Ippen, T., Mørk, H., Jordan, J., Senk, J., Konradi, S., Weidel, P., Fardet, T., Dahmen, D., Terhorst, D., Stapmanns, J., Trensch, G., … Plesser, H. E. (2021). NEST 3.0. Zenodo. 10.5281/ZENODO.4739103

27. Heiney, S. A., Kim, J., Augustine, G. J., & Medina, J. F. (2014). Precise Control of Movement Kinematics by Optogenetic Inhibition of Purkinje Cell Activity. Journal of Neuroscience, 34(6), 2321–2330. 10.1523/JNEUROSCI.4547-13.2014

28. Heinzle, J., König, P., & Salazar, R. F. (2007). Modulation of synchrony without changes in firing rates. Cognitive Neurodynamics 2007 1:3, 1(3), 225–235. 10.1007/S11571-007-9017-X

29. Kusch, L., Diaz-Pier, S., Klijn, W., Sontheimer, K., Bernard, C., Morrison, A., & Jirsa, V. (2024). Multiscale co-simulation design pattern for neuroscience applications. Frontiers in Neuroinformatics, 18, 1156683. 10.3389/FNINF.2024.1156683

30. Lindeman, S., Hong, S., Kros, L., Mejias, J. F., Romano, V., Oostenveld, R., Negrello, M., Bosman, L. W. J., & de Zeeuw, C. I. (2021). Cerebellar Purkinje cells can differentially modulate coherence between sensory and motor cortex depending on region and behavior. Proceedings of the National Academy of Sciences, 118(2), e2015292118. 10.1073/PNAS.2015292118

31. Lowet, E., Roberts, M. J., Bonizzi, P., Karel, J., & De Weerd, P. (2016). Quantifying Neural Oscillatory Synchronization: A Comparison between Spectral Coherence and Phase-Locking Value Approaches. PLOS ONE, 11(1), e0146443. 10.1371/JOURNAL.PONE.0146443

32. Marshall, J. D., Li, J. Z., Zhang, Y., Gong, Y., St-Pierre, F., Lin, M. Z., & Schnitzer, M. J. (2016). Cell-Type-Specific Optical Recording of Membrane Voltage Dynamics in Freely Moving Mice. Cell, 167(6), 1650–1662.e15. 10.1016/j.cell.2016.11.021

33. McAfee, S. S., Liu, Y., Sillitoe, R. V., & Heck, D. H. (2022). Cerebellar Coordination of Neuronal Communication in Cerebral Cortex. Frontiers in Systems Neuroscience, 15, 781527. 10.3389/FNSYS.2021.781527/FULL

34. Meier, J. M., Perdikis, D., Blickensdörfer, A., Stefanovski, L., Liu, Q., Maith, O., Dinkelbach, H. Ü., Baladron, J., Hamker, F. H., & Ritter, P. (2022). Virtual deep brain stimulation: Multiscale co-simulation of a spiking basal ganglia model and a whole-brain mean-field model with The Virtual Brain. Experimental Neurology, 354, 114111. 10.1016/j.expneurol.2022.114111

35. Melozzi, F., Woodman, M. M., Jirsa, V. K., & Bernard, C. (2017). The Virtual Mouse Brain: A Computational Neuroinformatics Platform to Study Whole Mouse Brain Dynamics. ENeuro, 4(3). 10.1523/ENEURO.0111-17.2017

36. Middleton, S. J., Racca, C., Cunningham, M. O., Traub, R. D., Monyer, H., Knöpfel, T., Schofield, I. S., Jenkins, A., & Whittington, M. A. (2008). High-Frequency Network Oscillations in Cerebellar Cortex. Neuron, 58(5), 763–774. 10.1016/j.neuron.2008.03.030

37. Ódor, G., Deco, G., & Kelling, J. (2022). Differences in the critical dynamics underlying the human and fruit-fly connectome. Physical Review Research, 4(2). 10.1103/PHYSREVRESEARCH.4.023057

38. Oh, S. W., Harris, J. A., Ng, L., Winslow, B., Cain, N., Mihalas, S., Wang, Q., Lau, C., Kuan, L., Henry, A. M., Mortrud, M. T., Ouellette, B., Nguyen, T. N., Sorensen, S. A., Slaughterbeck, C. R., Wakeman, W., Li, Y., Feng, D., Ho, A., … Zeng, H. (2014). A mesoscale connectome of the mouse brain. Nature, 508(7495), 207–214. 10.1038/nature13186

39. Palesi, F., Lorenzi, R. M., Casellato, C., Ritter, P., Jirsa, V., Gandini Wheeler-Kingshott, C. A. M., & D’Angelo, E. (2020). The Importance of Cerebellar Connectivity on Simulated Brain Dynamics. Frontiers in Cellular Neuroscience, 14, 570527. 10.3389/FNCEL.2020.00240/TEXT

40. Person, A. L., & Raman, I. M. (2011). Purkinje neuron synchrony elicits time-locked spiking in the cerebellar nuclei. Nature, 481(7382), 502–505. 10.1038/NATURE10732

41. Pisano, T. J., Dhanerawala, Z. M., Kislin, M., Bakshinskaya, D., Engel, E. A., Hansen, E. J., Hoag, A. T., Lee, J., de Oude, N. L., Venkataraju, K. U., Verpeut, J. L., Hoebeek, F. E., Richardson, B. D., Boele, H. J., & Wang, S. S. H. (2021). Homologous organization of cerebellar pathways to sensory, motor, and associative forebrain. Cell Reports, 36(12). 10.1016/j.celrep.2021.109721

42. Popa, D., Spolidoro, M., Proville, R. D., Guyon, N., Belliveau, L., & Léna, C. (2013). Functional role of the cerebellum in gamma-band synchronization of the sensory and motor cortices. The Journal of Neuroscience, 33(15), 6552–6556. 10.1523/JNEUROSCI.5521-12.2013

43. Proville, R. D., Spolidoro, M., Guyon, N., Dugué, G. P., Selimi, F., Isope, P., Popa, D., & Léna, C. (2014). Cerebellum involvement in cortical sensorimotor circuits for the control of voluntary movements. Nature Neuroscience, 17(9), 1233–1239. 10.1038/NN.3773;TECHMETA

44. Rancz, E. A., Ishikawa, T., Duguid, I., Chadderton, P., Mahon, S., & Häusser, M. (2007). High-fidelity transmission of sensory information by single cerebellar mossy fibre boutons. Nature, 450(7173), 1245–1248. 10.1038/nature05995

45. Ritter, P., Schirner, M., Mcintosh, A. R., & Jirsa, V. K. (2013). The virtual brain integrates computational modeling and multimodal neuroimaging. Brain Connectivity, 3(2), 121–145. 10.1089/BRAIN.2012.0120

46. Riva, D., Annunziata, S., Contarino, V., Erbetta, A., Aquino, D., & Bulgheroni, S. (2013). Gray Matter Reduction in the Vermis and CRUS-II Is Associated with Social and Interaction Deficits in Low-Functioning Children with Autistic Spectrum Disorders: a VBM-DARTEL Study. The Cerebellum 2013 12:5, 12(5), 676–685. 10.1007/S12311-013-0469-8

47. Robinson, P. A., Rennie, C. J., Wright, J. J., Bahramali, H., Gordon, E., & Rowe, D. L. (2001). Prediction of electroencephalographic spectra from neurophysiology. Physical Review. E, Statistical, Nonlinear, and Soft Matter Physics, *63*(2 Pt 1), 021903. 10.1103/PhysRevE.63.021903

48. Rocha, R. P., Koçillari, L., Suweis, S., Corbetta, M., & Maritan, A. (2018). Homeostatic plasticity and emergence of functional networks in a whole-brain model at criticality. Scientific Reports 2018 8:1, 8(1), 15682-. 10.1038/s41598-018-33923-9

49. Romano, V., de Propris, L., Bosman, L. W. J., Warnaar, P., Ten Brinke, M. M., Lindeman, S., Ju, C., Velauthapillai, A., Spanke, J. K., Guerra, E. M., Hoogland, T. M., Negrello, M., D’angelo, E., & De Zeeuw, C. I. (2018). Potentiation of cerebellar purkinje cells facilitates whisker reflex adaptation through increased simple spike activity. ELife, 7. 10.7554/ELIFE.38852

50. Romano, V., Reddington, A. L., Cazzanelli, S., Mazza, R., Ma, Y., Strydis, C., Negrello, M., Bosman, L. W. J., & De Zeeuw, C. I. (2020). Functional Convergence of Autonomic and Sensorimotor Processing in the Lateral Cerebellum. Cell Reports, 32(1), 107867. 10.1016/J.CELREP.2020.107867

51. Sanz-Leon, P., Knock, S. A., Spiegler, A., & Jirsa, V. K. (2015). Mathematical framework for large-scale brain network modeling in The Virtual Brain. NeuroImage, 111, 385–430. 10.1016/J.NEUROIMAGE.2015.01.002

52. Sanz-Leon, P., Knock, S. A., Woodman, M. M., Domide, L., Mersmann, J., Mcintosh, A. R., & Jirsa, V. (2013). The virtual brain: A simulator of primate brain network dynamics. Frontiers in Neuroinformatics, 7(MAY), 47900. 10.3389/FNINF.2013.00010/ABSTRACT

53. Schirner, M., Deco, G., & Ritter, P. (2023). Learning how network structure shapes decision-making for bio-inspired computing. Nature Communications 2023 14:1, 14(1), 2963-. 10.1038/s41467-023-38626-y

54. Schirner, M., Domide, L., Perdikis, D., Triebkorn, P., Stefanovski, L., Pai, R., Prodan, P., Valean, B., Palmer, J., Langford, C., Blickensdörfer, A., van der Vlag, M., Diaz-Pier, S., Peyser, A., Klijn, W., Pleiter, D., Nahm, A., Schmid, O., Woodman, M., … Ritter, P. (2022). Brain simulation as a cloud service: The Virtual Brain on EBRAINS. NeuroImage, 251. 10.1016/j.neuroimage.2022.118973

55. Schirner, M., McIntosh, A. R., Jirsa, V., Deco, G., & Ritter, P. (2018). Inferring multi-scale neural mechanisms with brain network modelling. ELife, 7. 10.7554/eLife.28927

56. Shin, S. L., & De Schutter, E. (2006). Dynamic Synchronization of Purkinje Cell Simple Spikes. 10.1152/Jn.00570.2006, 96(6), 3485–3491. 10.1152/JN.00570.2006

57. Sober, S. J., & Sabes, P. N. (2005). Flexible strategies for sensory integration during motor planning. Nature Neuroscience, 8(4), 490. 10.1038/NN1427

58. Sreenivasan, V., Esmaeili, V., Kiritani, T., Galan, K., Crochet, S., & Petersen, C. C. H. (2016). Movement initiation signals in mouse whisker motor cortex. Neuron, 92(6), 1368–1382. 10.1016/j.neuron.2016.12.001

59. Stasinski, J., Taher, H., Meier, J. M., Schirner, M., Perdikis, D., & Ritter, P. (2024). Homeodynamic feedback inhibition control in whole-brain simulations. PLOS Computational Biology, 20(12), e1012595. 10.1371/JOURNAL.PCBI.1012595

60. Tejero-Cantero, A., Boelts, J., Deistler, M., Lueckmann, J.-M., Durkan, C., Gonçalves, P., Greenberg, D., & Macke, J. (2020). sbi: A toolkit for simulation-based inference. Journal of Open Source Software, 5(52), 2505. 10.21105/JOSS.02505/STATUS.SVG)

61. ten Brinke, M. M., Boele, H. J., Spanke, J. K., Potters, J. W., Kornysheva, K., Wulff, P., IJpelaar, A. C. H. G., Koekkoek, S. K. E., & De Zeeuw, C. I. (2015). Evolving Models of Pavlovian Conditioning: Cerebellar Cortical Dynamics in Awake Behaving Mice. Cell Reports, 13(9), 1977–1988. 10.1016/j.celrep.2015.10.057

62. ten Brinke, M. M., Heiney, S. A., Wang, X., Proietti-Onori, M., Boele, H. J., Bakermans, J., Medina, J. F., Gao, Z., & De Zeeuw, C. I. (2017). Dynamic modulation of activity in cerebellar nuclei neurons during pavlovian eyeblink conditioning in mice. ELife, 6, 1–27. 10.7554/eLife.28132

63. Timmann, D., Watts, S., & Hore, J. (1999). Failure of cerebellar patients to time finger opening precisely causes ball high-low inaccuracy in overarm throws. Journal of Neurophysiology, 82(1), 103–114. 10.1152/JN.1999.82.1.103

64. Triebkorn, P., Meier, J., Zimmermann, J., Stefanovski, L., Roy, D., Solodkin, A., Jirsa, V., Deco, G., Breakspear, M., Schirner, M., McIntosh, A. R., & Ritter, P. (2024). Fifty shades of The Virtual Brain: Converging optimal working points yield biologically plausible electrophysiological and imaging features. BioRxiv, 2020.03.26.009795. 10.1101/2020.03.26.009795

65. van der Heijden, M. E., Brown, A. M., & Sillitoe, R. V. (2022). Influence of data sampling methods on the representation of neural spiking activity in vivo. IScience, 25(11). 10.1016/j.isci.2022.105429

66. Van Der Werf, Y. D., Witter, M. P., & Groenewegen, H. J. (2002). The intralaminar and midline nuclei of the thalamus. Anatomical and functional evidence for participation in processes of arousal and awareness. Brain Research Reviews, 39(2–3), 107–140. 10.1016/S0165-0173(02)00181-9

67. White, E. L. (1979). Thalamocortical synaptic relations: A review with emphasis on the projections of specific thalamic nuclei to the primary sensory areas of the neocortex. Brain Research Reviews, 1(3), 275–311. 10.1016/0165-0173(79)90008-0

68. Wolpert, D. M., Ghahramani, Z., & Jordan, M. I. (1995). An internal model for sensorimotor integration. Science, 269(5232), 1880–1882. 10.1126/SCIENCE.7569931;PAGE:STRING:ARTICLE/CHAPTER

69. Wu, S., Wardak, A., Khan, M. M., Chen, C. H., & Regehr, W. G. (2024). Implications of variable synaptic weights for rate and temporal coding of cerebellar outputs. ELife, 13. 10.7554/ELIFE.89095

70. Wu, Y., & Raman, I. M. (2017). Facilitation of mossy fibre-driven spiking in the cerebellar nuclei by the synchrony of inhibition. The Journal of Physiology, 595(15), 5245. 10.1113/JP274321

