## Supplementary Figures, Tables and Notes for "The Cerebellar Engine: Multiscale Digital Brain Co-simulations Reveal How Cerebellar Spiking Architecture Shapes Cortical Coherence"

### Supplementary Information

Alice Geminiani[1,*,+] Jil Mona Meier[2,3,*,+], Dionysios Perdikis[2,3,*,+], Sabrine Ouertani[2,3], Claudia Casellato[1], Petra Ritter[2,3,4,5,6,T] , Egidio D'Angelo[1,7,T]

[1] Neurophysiology Unit, Dept of Brain and Behavioral Sciences, University of Pavia, Pavia, Italy
[2] Berlin Institute of Health at Charité – Universitätsmedizin Berlin, Charitéplatz 1, 10117, Berlin, Germany
[3] Department of Neurology with Experimental Neurology, Charité – Universitätsmedizin Berlin, corporate member of Freie Universität Berlin and Humboldt-Universität zu Berlin, Charitéplatz 1, 10117, Berlin, Germany
[4] Bernstein Focus State Dependencies of Learning and Bernstein Center for Computational Neuroscience, Berlin, Germany
[5] Einstein Center for Neuroscience Berlin, Charitéplatz 1, 10117, Berlin, Germany
[6] Einstein Center Digital Future, Wilhelmstraße 67, 10117, Berlin, Germany
[7] Digital Neuroscience Center, IRCCS Mondino Foundation, Pavia, Italy

[*,T] equal contribution

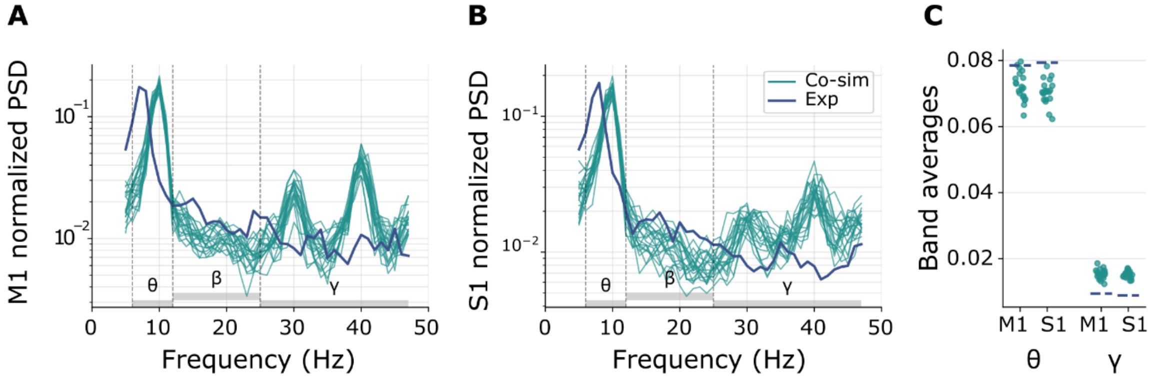


***Supplementary Figure S1. Oscillation power of M1 and S1 in TVMB-NEST co-simulations.*** *PSD of* ***(A)*** *M1 and* ***(B)*** *S1 for 10 TVMB-NEST co-simulation runs and both hemispheres (n=20) represented in green, compared to the target experimental PSD from (Popa et al., 2013) in blue.* ***(C)*** *Average power in theta and gamma band.*

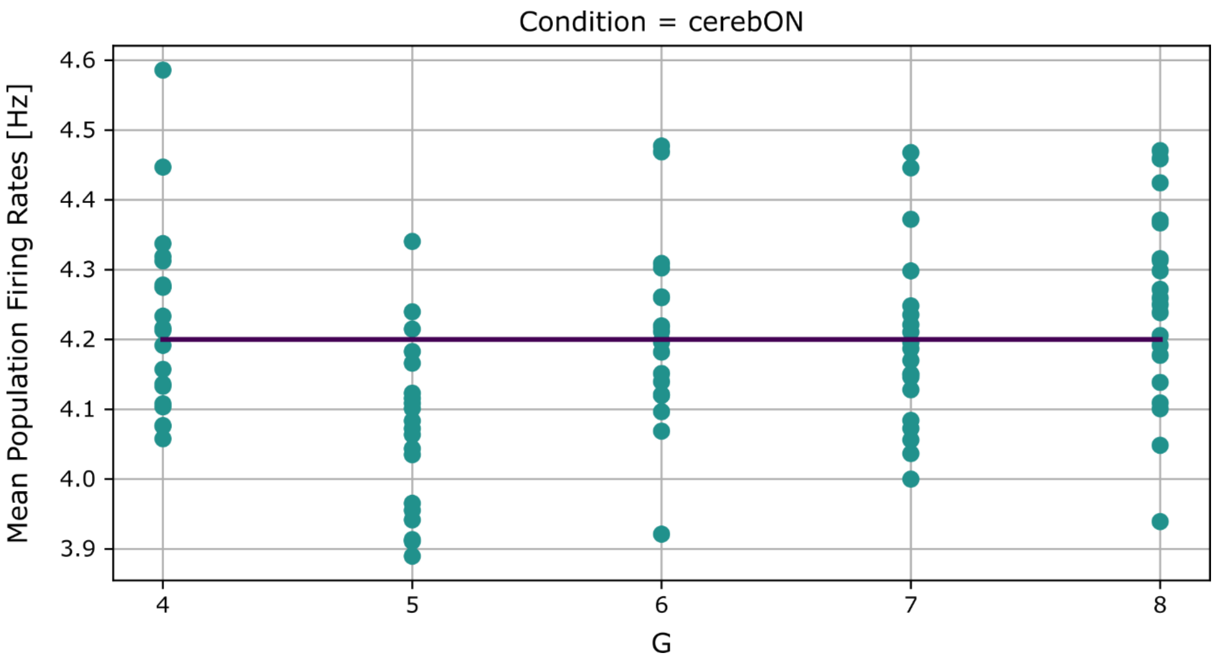


***Supplementary Figure S2. Ansiform Lobule Granule Cells (GrC) average population firing rate for validation of tuning of TVMB-NEST parameters for a range of values of G via co-simulations.*** *Average GrC population firing rates from 10 repetitions of TVMB-NEST co-simulations with the fitted parameters are depicted for both hemispheres in teal filled circles for each G value in the range of [4, 8] shown as x tick labels. The target firing rate of 4.2 spikes/s is shown as a deep purple line.*

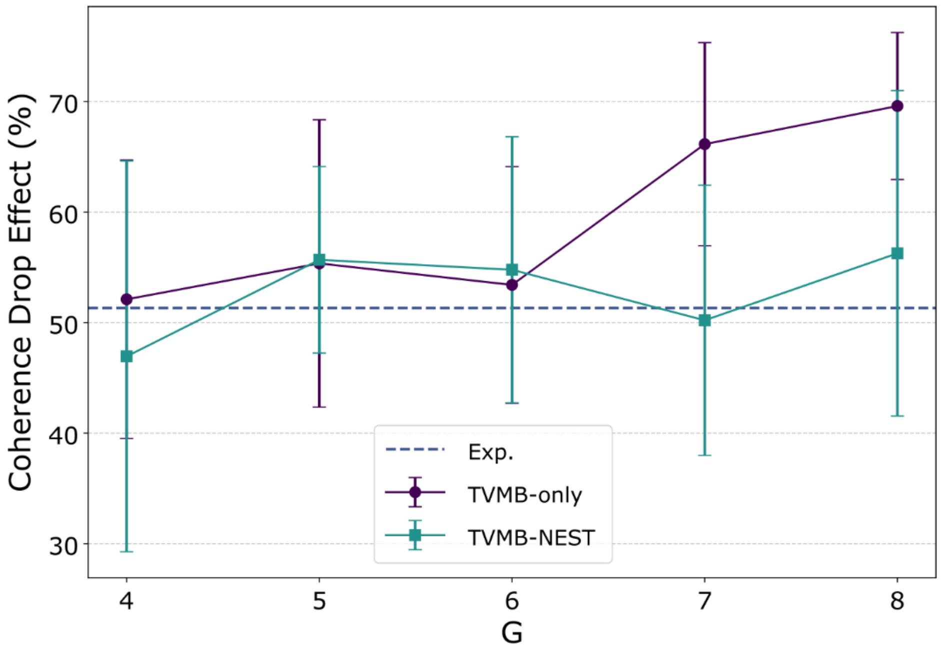


***Supplementary Figure S3. Coherence drop for different G values.*** *M1-S1 coherence drop effect as percentage of the normalized difference (cerebON – cerebOFF) / cerebON in gamma band (25 – 60 Hz) when inactivating the cerebellum, in TVMB-only simulations and TVMB-NEST co-simulations for all G values, displayed as means and standard deviation (over the pooled hemispheric values and 10 simulation runs for each hemisphere). The dashed blue horizontal line displays the experimental data taken from (Popa et al., 2013), where mean gamma-band coherence drops between M1 and S1 were similarly calculated for comparison.*

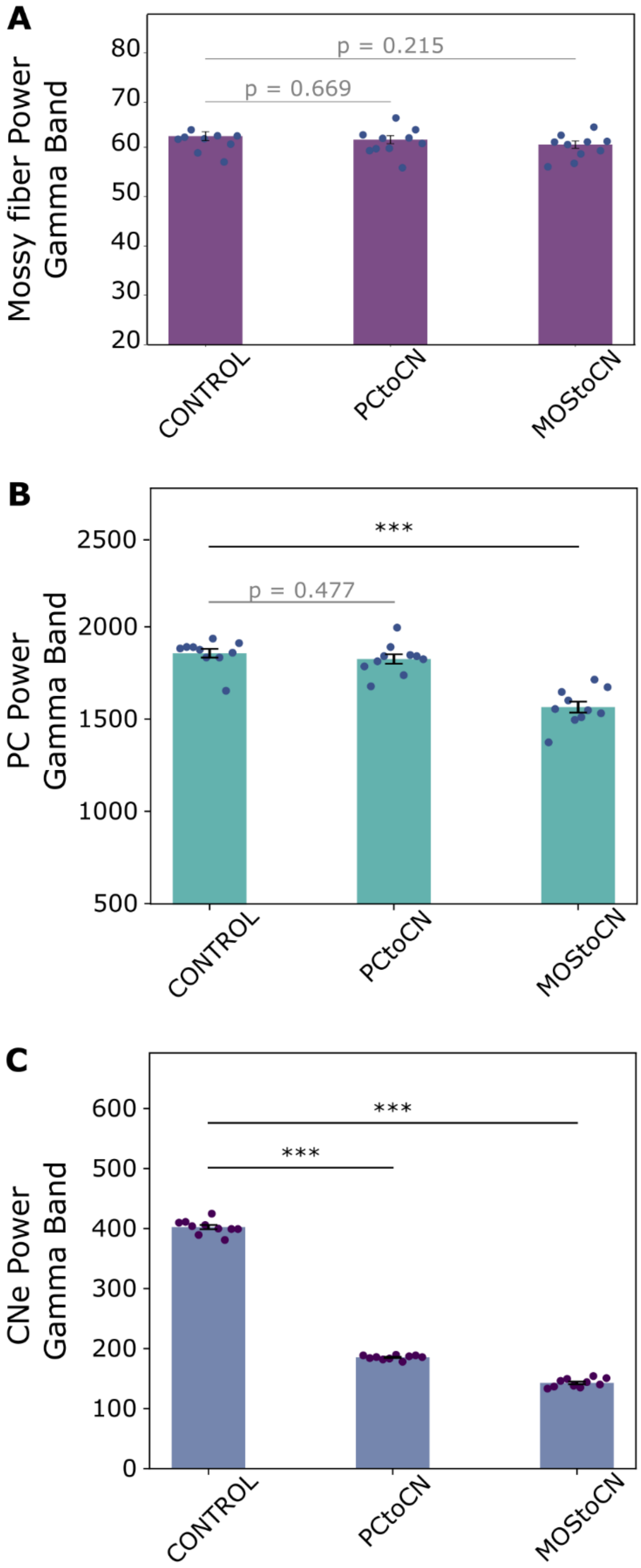


***Supplementary Figure S4. Gamma-band power inside the cerebellar circuit.*** *Bar plot of the total power in gamma band for* ***(A)*** *mossy fibers,* ***(B)*** *Purkinje cells, PC and* ***(C)*** *excitatory cerebellar nuclei neurons, CNe in CONTROL and in virtual lesion conditions – PCtoCN and MOStoCN. (n = 10 independent samples, averaged over both hemispheres, mean ± STD, ***p < 0.001; two-sided unpaired t-test). PCtoCN: inactivating the indirect inhibitory projection from PC to CNe; MOStoCN: inactivating the direct excitatory projection from mossy fibers to CNe; CNe: excitatory cerebellar nuclei neurons.*

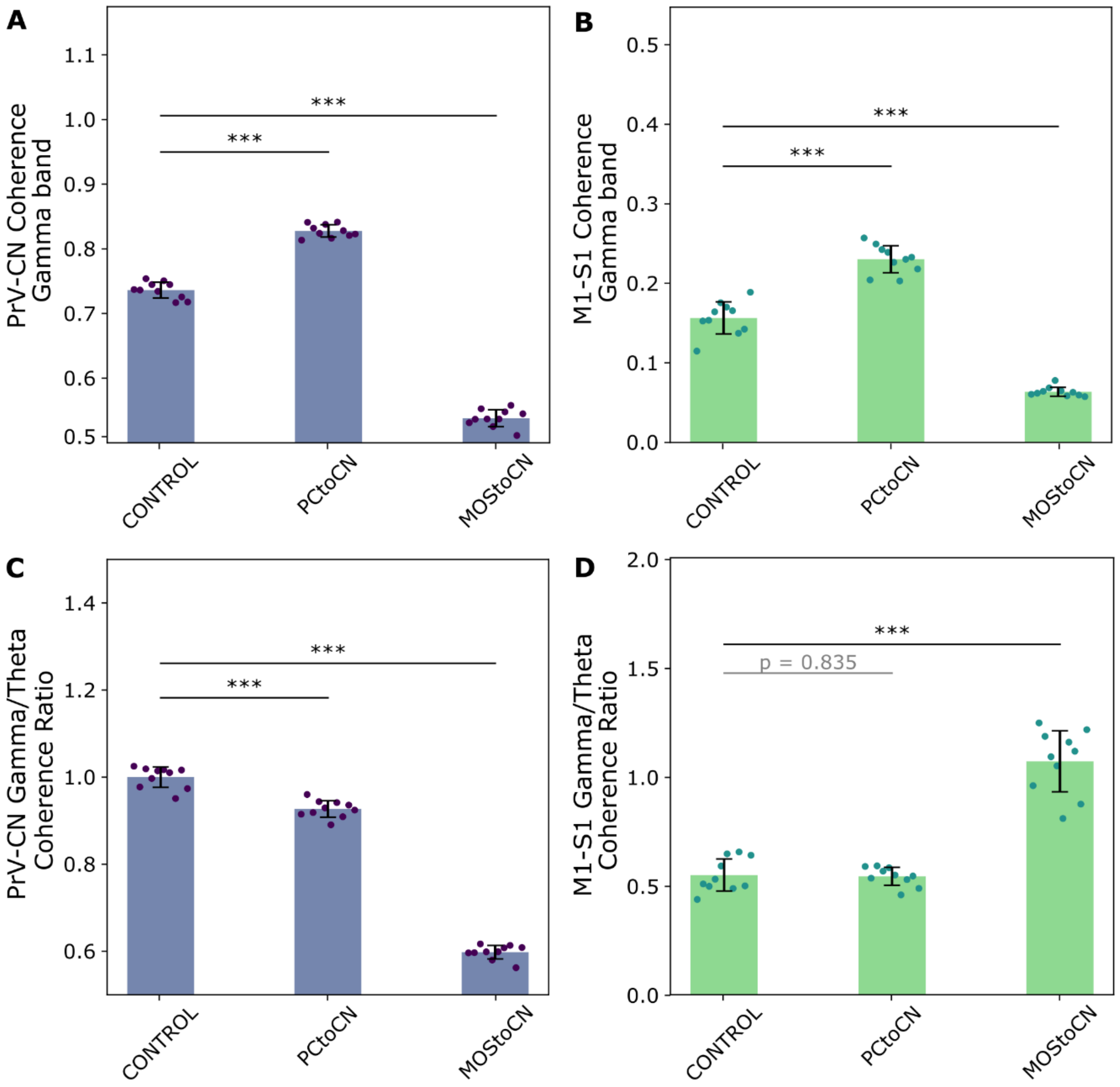


***Supplementary Figure S5. Coherence in non-normalized cerebellar output simulations. (A)*** *Gamma-band for cerebellar input-output and* ***(B)*** *M1-S1, in CONTROL and simulations with separate ablation of CN inputs with non-normalized cerebellar output.* ***(C)*** *Gamma-to-theta coherence for cerebellar input-output and* ***(D)*** *M1-S1 in the same conditions. Bar plots and error bars respectively show the mean and standard deviation over 10 simulations, each one averaged across the two hemispheres, with comparisons between CONTROL and each lesion condition (*p < 0.05, **p < 0.01; independent-samples two-sided unpaired t-test, FDR-corrected over all lesion conditions).*

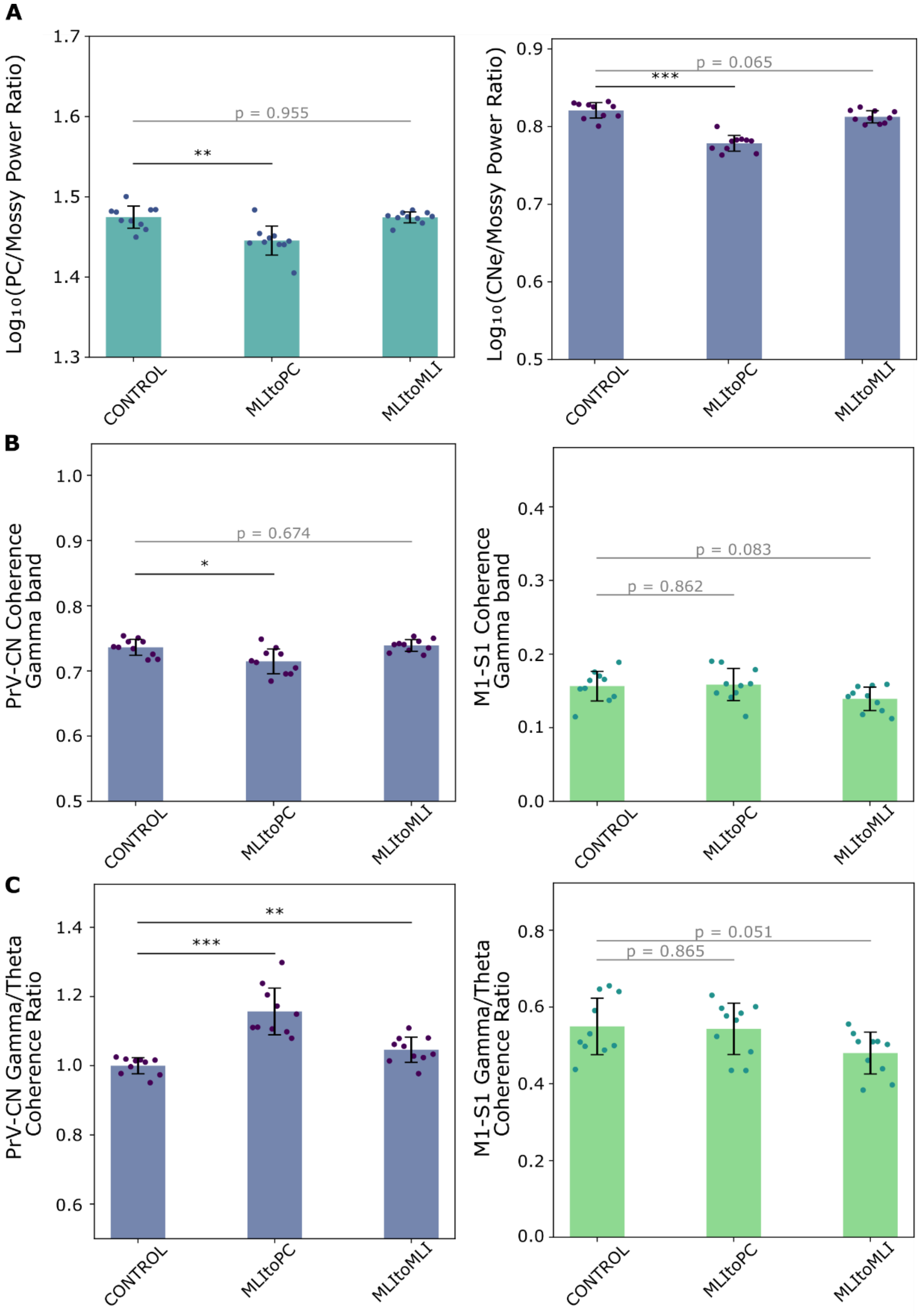


***Supplementary Figure S6. Gamma power and coherence with MLI lesions. (A)*** *Relative change of power in two output populations (Purkinje-left and CN-right) with respect to mossy fibers (power ratio) in CONTROL and MLI lesions (n = 10 independent samples, mean ± STD, *p < 0.05, **p < 0.01, ***p < 0.001; two-sided unpaired t-test).* ***(B)*** *Gamma-band and* ***(C)*** *gamma-to-theta coherence for PrV-CN (cerebellar input-output) and M1-S1, in CONTROL and simulations with separate ablation of inhibition to PC and self-inhibition in the molecular layer. Bar plots and error bars respectively show the mean and standard deviation over 10 simulations, each one averaged*

*across the two hemispheres, with comparisons between CONTROL and each lesion condition (**p < 0.01; independent-samples two-sided unpaired t-test, FDR-corrected over all lesion conditions).*

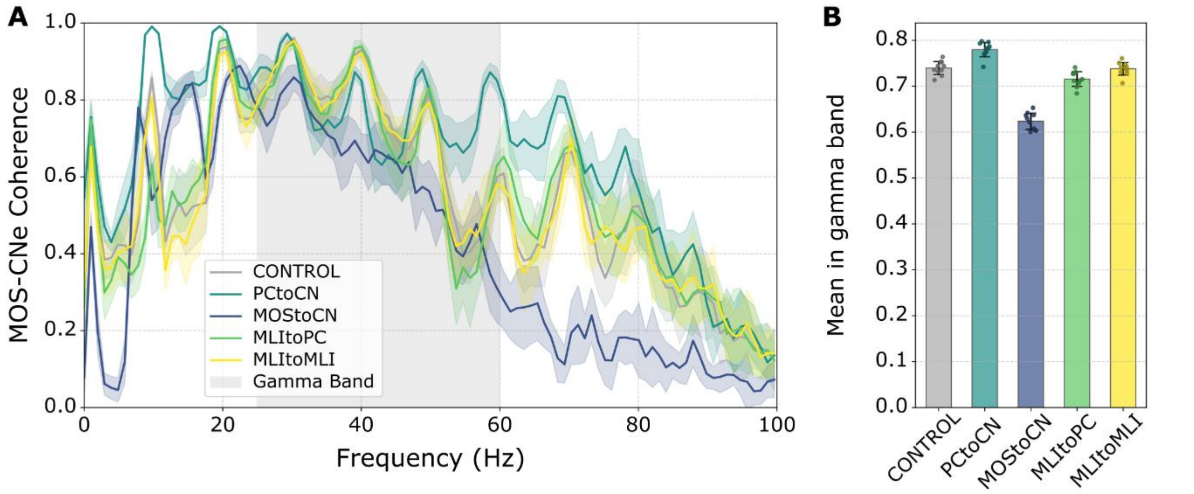


***Supplementary Figure S7. Coherence between mossy fibers and CNe population activity. (A)*** *Spectral coherence for CONTROL and all lesion conditions, represented as mean ± STD of 10 simulation runs for each condition, averaging over hemispheres. The gamma band is highlighted as the grey area.* ***(B)*** *Average coherence in the gamma band with comparisons between CONTROL and each lesion condition (***p < 0.001; independent-samples two-sided unpaired t-test, FDR-corrected over all lesion conditions).*

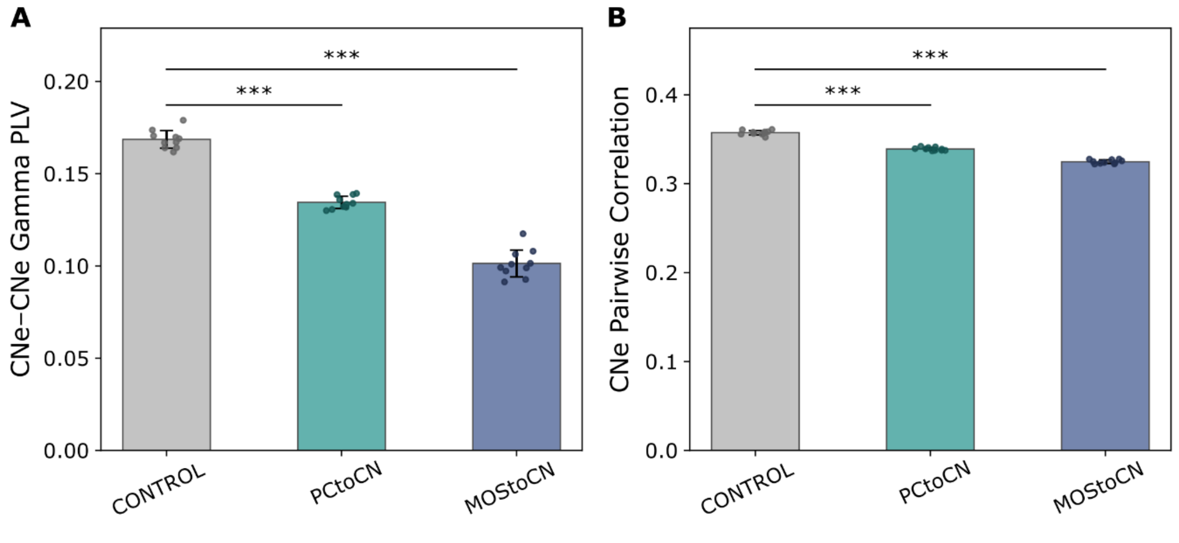


***Supplementary Figure S8. Intra-population spiking synchronization measures. (A)*** *PLV of CNe spikes relative to CNe population firing rate in the gamma band,* ***(B)*** *CNe population synchronization as pairwise spike train correlation in time bins of 10 ms. For each condition, mean ± STD across 10 simulation runs, averaged across hemispheres, is represented as a bar plot with errorbar. In all panels, the comparisons between CONTROL and each lesion condition are reported (***p<0.001; independent-samples two-sided unpaired t-test, FDR-corrected over all lesion conditions).*

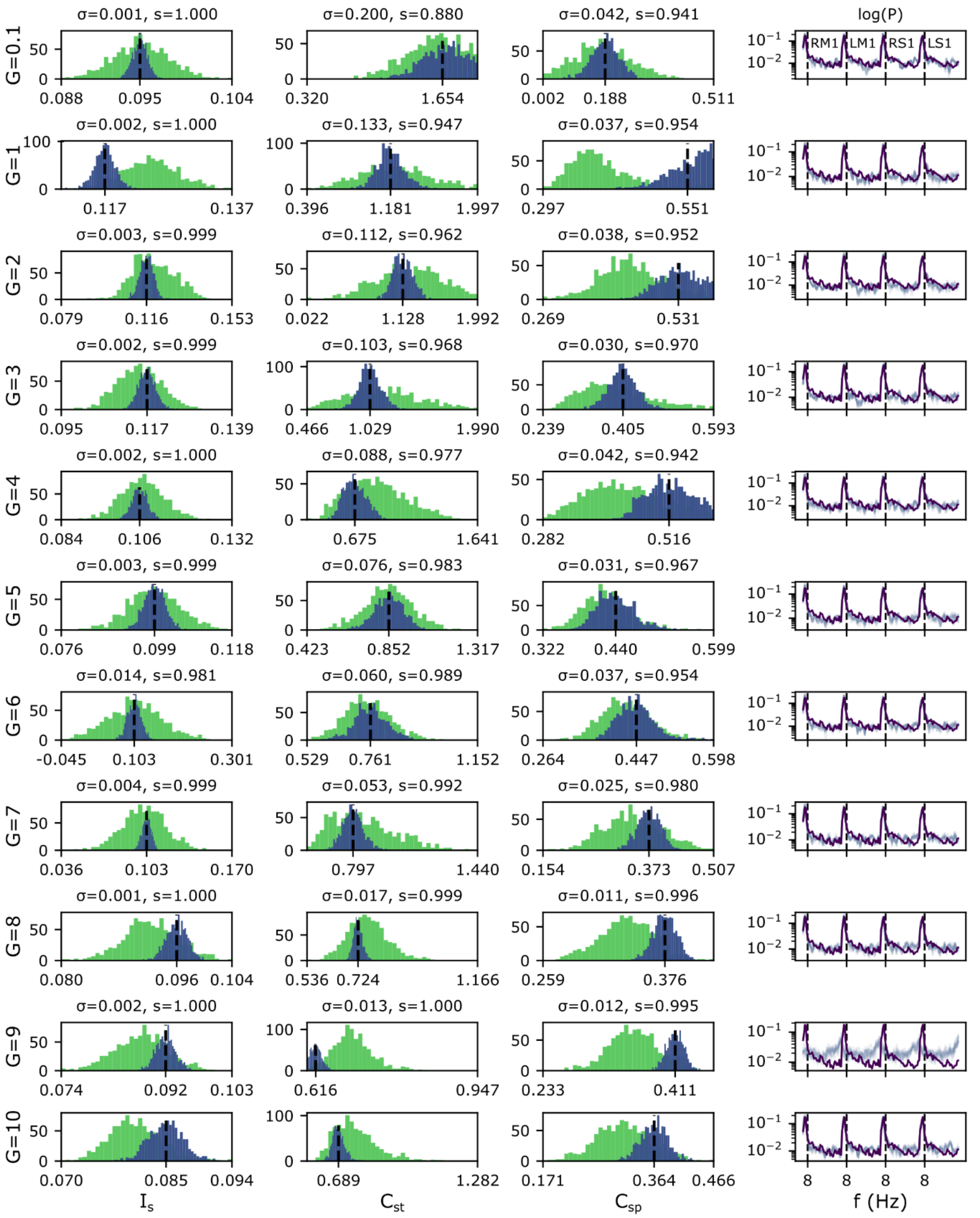


***Supplementary Figure S9. Fitting of parameters $I_S$, $C_{st}$ and $C_{sp}$ for a range of values of G.*** *Results of fitting parameters parameter $I_S$ (thalamic relay nucleus constant input in mV) and dimensionless scaling parameters $C_{st}$ (strength of scaling) and $C_{sp}$ (splitting between $C_{I_E}$ and $C_{W_{IE}}$, which scale per region connectivity indegree), for different values of global connectivity scaling factor (global coupling) G, via TVMB-only simulations without any whisking-task-related parameters (all values are rounded to 3 decimal points for display, see also Supplementary Note S1, Supplementary Table S4 and Methods 4.2.2 of the main manuscript). Each row corresponds to a value of G shown as labels of the first column from the left. Columns 1 to 3, starting from the left to the right, correspond to the three fitted parameters ($I_S$, $C_{st}$ and $C_{sp}$), as shown in the titles of the first row and the x labels of the last row starting from top to bottom. For each G value (rows) and parameter (columns) two histograms are depicted: the posterior distribution of the first round of fitting, acting also as prior distribution for the second round of fitting (in*

light green) and the final, posterior distribution of round 2 (in blue purple). The mean value of the parameters used for further simulations are shown by the black dotted line and the exact value is also shown as a tick label of the x axis. The standard deviation (σ) and shrinkage statistical value (s) between the prior distribution of the first round and the final, posterior distribution, are shown in the title of each panel. The fourth column depicts the power spectra resulting from 10 simulations using the mean values of the parameters for each G value (row) in faded blue purple color, as well as the target power spectra taken from (Popa et al., 2013) in deep purple, with a logarithmic y axis. The power spectra of Right M1, Left M1, Right S1 and Left S1 are concatenated from left to right as the labels show at the first row from top to bottom. We observe that each round of fitting led to higher accuracy and a narrower posterior distribution, whereas the target and the resulting power spectra match well, except for the case of G=9.0. Eventually, based on the above results, G values from 4 to 8 were used for further simulations, i.e., excluding values of low G (which would weaken all interactions among brain regions) and higher G values, starting from G=9.0, for which the fitting was not equally successful.

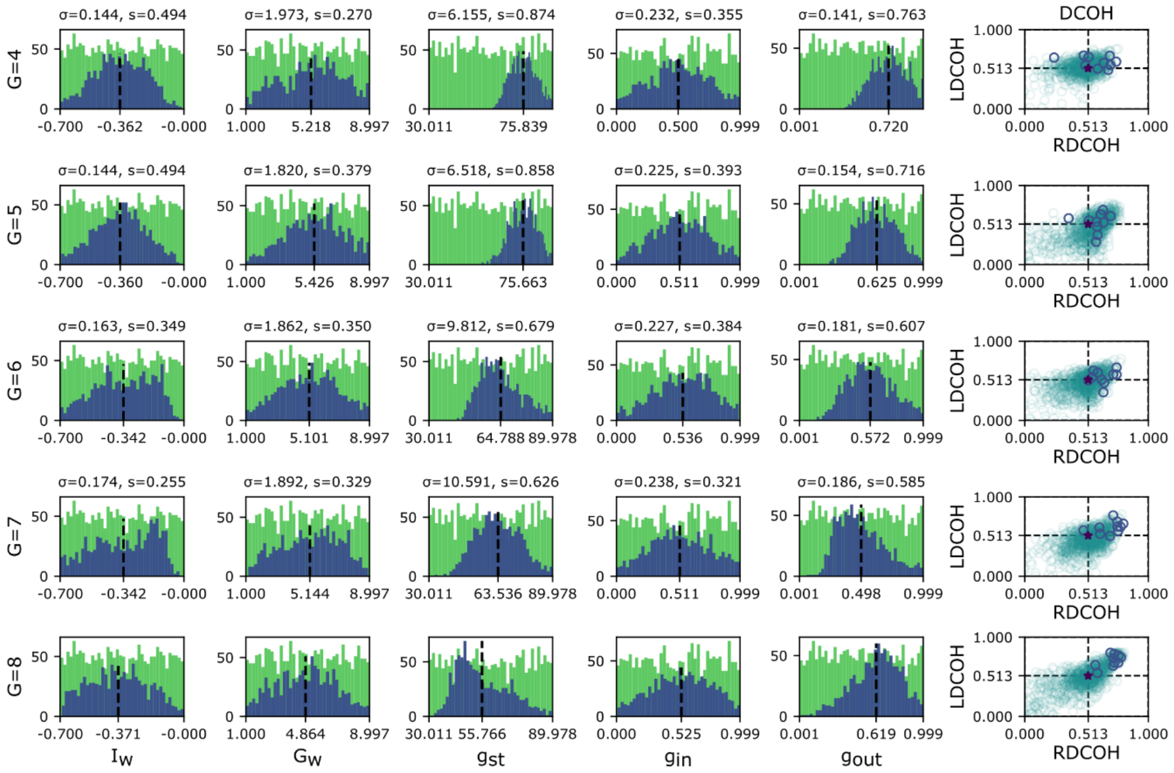


***Supplementary Figure S10. Fitting of whisking-task parameters for a range of G values via TVMB-only simulations.*** *Parameters include $I_W$ (whisking nodes' constant input in mV), $G_W$ (whisking nodes' long-range coupling scaling parameter,) $g_{st}$ (scaling the strength of all pathway gains), $g_{in}$ (scaling specific to the gain of the connection from M1 to FMN), $g_{out}$ (scaling specific to the gain of the connection from CN to the specific thalamic nucleus of M1), all scaling parameters being dimensionless (all values are rounded to 3 decimal points for display; see also Supplementary Note S1, Supplementary Table S5 and Methods 4.2.2 of the main manuscript). Each row corresponds to a value of G shown as labels of the first column from the left. Columns 1 to 5, starting from the left to the right, correspond to the five fitted parameters $I_W$, $G_W$, $g_{st}$, $g_{in}$ and $g_{out}$), as shown in the titles of the first row and the x labels of the last row starting from top to bottom. For each G value (rows) and parameter (columns) two histograms are depicted: the prior uniform distributions (not fully shown) in light green and the posterior distributions in blue purple. The mean value of the parameters used for further simulations are shown by the black dotted line and the exact value is also shown as a tick label of the x axis. The standard deviation (σ) and shrinkage statistical value (s) are shown in the title of each panel. The sixth column depicts: (a) the target DCOH, measure, i.e., the normalized gamma-band coherence drop between M1 and S1 (right at x axis -RDCOH x axis label- and left at y axis -LDCOH y axis label), after cerebellum deactivation, computed from data extracted from (Popa et al., 2013), as a deep purple asterisk, (b) the value of the same measure from 10 repetitions of TVMB-only simulations using the mean values of the fitted parameters in blue purple unfilled circles, and (c) the same measure for the values of the parameters of the 1,000 samples of the posterior distribution, averaged of three repetitions, in faded, unfilled, teal circles. The high shrinkage values and the proximity of the blue purple circles (from simulations with mean values) as well as most of the teal circles (from samples of the posterior distribution) shows that the fitting was*

*successful for all G values, whereas the range covered by the posterior distributions, considered together with the range covered by the green circles in the space of the measure, provides some evidence on the - relatively low - sensitivity of the result. In other words, the accuracy of the fitting is not crucial for further simulations and the results of the current study.*

***Supplementary Table S1: Region labels and corresponding major region labels from the 298 regions from the Allen mouse brain atlas connectome generation.*** *The extracted connectome (before merging) consisted of a respective right and left hemispheric region of each of these regions (596 regions in total).*

| Region Name | Major Region |
|---|---|
| Frontal pole, cerebral cortex | Isocortex |
| Primary motor area | Isocortex |
| Secondary motor area | Isocortex |
| Primary somatosensory area, nose | Isocortex |
| Primary somatosensory area, barrel field | Isocortex |
| Primary somatosensory area, lower limb | Isocortex |
| Primary somatosensory area, mouth | Isocortex |
| Primary somatosensory area, upper limb | Isocortex |
| Primary somatosensory area, trunk | Isocortex |
| Primary somatosensory area, unassigned | Isocortex |
| Supplemental somatosensory area | Isocortex |
| Gustatory areas | Isocortex |
| Visceral area | Isocortex |
| Dorsal auditory area | Isocortex |
| Primary auditory area | Isocortex |
| Posterior auditory area | Isocortex |
| Ventral auditory area | Isocortex |
| Anterolateral visual area | Isocortex |
| Anteromedial visual area | Isocortex |
| Lateral visual area | Isocortex |
| Primary visual area | Isocortex |
| Posterolateral visual area | Isocortex |
| posteromedial visual area | Isocortex |
| Laterointermediate area | Isocortex |
| Postrhinal area | Isocortex |
| Anterior cingulate area, dorsal part | Isocortex |
| Anterior cingulate area, ventral part | Isocortex |
| Prelimbic area | Isocortex |
| Infralimbic area | Isocortex |
| Orbital area, lateral part | Isocortex |
| Orbital area, medial part | Isocortex |
| Orbital area, ventrolateral part | Isocortex |
| Agranular insular area, dorsal part | Isocortex |
| Agranular insular area, posterior part | Isocortex |
| Agranular insular area, ventral part | Isocortex |
| Retrosplenial area, lateral agranular part | Isocortex |
| Retrosplenial area, dorsal part | Isocortex |
| Retrosplenial area, ventral part | Isocortex |
| Anterior area | Isocortex |
| Rostrolateral visual area | Isocortex |
| Temporal association areas | Isocortex |
| Perirhinal area | Isocortex |
| Ectorhinal area | Isocortex |
| Main olfactory bulb | Olfactory Areas |
| Accessory olfactory bulb | Olfactory Areas |
| Anterior olfactory nucleus | Olfactory Areas |
| Taenia tecta | Olfactory Areas |
| Dorsal peduncular area | Olfactory Areas |
| Piriform area | Olfactory Areas |
| Nucleus of the lateral olfactory tract | Olfactory Areas |
| Cortical amygdalar area, anterior part | Olfactory Areas |
| Cortical amygdalar area, posterior part | Olfactory Areas |
| Piriform-amygdalar area | Olfactory Areas |
| Postpiriform transition area | Olfactory Areas |
| Field CA1 | Hippocampal Formation |
| Field CA2 | Hippocampal Formation |
| Field CA3 | Hippocampal Formation |
| Dentate gyrus | Hippocampal Formation |
| Induseum griseum | Hippocampal Formation |
| Entorhinal area, lateral part | Hippocampal Formation |
| Entorhinal area, medial part, dorsal zone | Hippocampal Formation |
| Parasubiculum | Hippocampal Formation |
| Postsubiculum | Hippocampal Formation |
| Presubiculum | Hippocampal Formation |
| Subiculum | Hippocampal Formation |
| Prosubiculum | Hippocampal Formation |

| | |
|---|---|
| Area prostriata | Isocortex |
| Claustrum | Cortical Subplate |
| Endopiriform nucleus, dorsal part | Cortical Subplate |
| Endopiriform nucleus, ventral part | Cortical Subplate |
| Lateral amygdalar nucleus | Cortical Subplate |
| Basolateral amygdalar nucleus | Cortical Subplate |
| Basomedial amygdalar nucleus | Cortical Subplate |
| Posterior amygdalar nucleus | Cortical Subplate |
| Caudoputamen | Striatum |
| Nucleus accumbens | Striatum |
| Fundus of striatum | Striatum |
| Olfactory tubercle | Striatum |
| Lateral septal nucleus, caudal (caudodorsal) part | Pallidum |
| Lateral septal nucleus, rostral (rostroventral) part | Pallidum |
| Lateral septal nucleus, ventral part | Pallidum |
| Septofimbrial nucleus | Pallidum |
| Anterior amygdalar area | Striatum |
| Bed nucleus of the accessory olfactory tract | Pallidum |
| Central amygdalar nucleus | Striatum |
| Intercalated amygdalar nucleus | Striatum |
| Medial amygdalar nucleus | Striatum |
| Globus pallidus, external segment | Pallidum |
| Globus pallidus, internal segment | Pallidum |
| Substantia innominata | Pallidum |
| Magnocellular nucleus | Pallidum |
| Medial septal nucleus | Pallidum |
| Diagonal band nucleus | Pallidum |
| Triangular nucleus of septum | Pallidum |
| Bed nuclei of the stria terminalis | Pallidum |
| Bed nucleus of the anterior commissure | Pallidum |
| Ventral anterior-lateral complex of the thalamus | Thalamus |
| Ventral medial nucleus of the thalamus | Thalamus |
| Ventral posterolateral nucleus of the thalamus | Thalamus |
| Ventral posterolateral nucleus of the thalamus, parvicellular part | Thalamus |
| Ventral posteromedial nucleus of the thalamus | Thalamus |
| Ventral posteromedial nucleus of the thalamus, parvicellular part | Thalamus |
| Posterior triangular thalamic nucleus | Thalamus |
| Subparafascicular nucleus, magnocellular part | Thalamus |
| Subparafascicular nucleus, parvicellular part | Thalamus |
| Subparafascicular area | Thalamus |
| Peripeduncular nucleus | Thalamus |
| Medial geniculate complex | Thalamus |
| Dorsal part of the lateral geniculate complex | Thalamus |
| Lateral posterior nucleus of the thalamus | Thalamus |
| Posterior complex of the thalamus | Thalamus |
| Posterior limiting nucleus of the thalamus | Thalamus |
| Suprageniculate nucleus | Thalamus |
| Anteroventral nucleus of thalamus | Thalamus |
| Anteromedial nucleus | Thalamus |
| Anterodorsal nucleus | Thalamus |
| Interanteromedial nucleus of the thalamus | Thalamus |
| Interanterodorsal nucleus of the thalamus | Thalamus |
| Lateral dorsal nucleus of thalamus | Thalamus |
| Intermediodorsal nucleus of the thalamus | Nonspecific Thalamus |
| Mediodorsal nucleus of thalamus | Thalamus |
| Submedial nucleus of the thalamus | Thalamus |
| Perireunensis nucleus | Nonspecific Thalamus |
| Paraventricular nucleus of the thalamus | Nonspecific Thalamus |
| Parataenial nucleus | Nonspecific Thalamus |
| Nucleus of reuniens | Nonspecific Thalamus |
| Xiphoid thalamic nucleus | Nonspecific Thalamus |
| Rhomboid nucleus | Nonspecific Thalamus |
| Central medial nucleus of the thalamus | Nonspecific Thalamus |
| Paracentral nucleus | Nonspecific Thalamus |
| Central lateral nucleus of the thalamus | Nonspecific Thalamus |
| Parafascicular nucleus | Nonspecific Thalamus |
| Posterior intralaminar thalamic nucleus | Nonspecific Thalamus |
| Reticular nucleus of the thalamus | Nonspecific Thalamus |
| Intergeniculate leaflet of the lateral geniculate complex | Thalamus |
| Intermediate geniculate nucleus | Thalamus |
| Ventral part of the lateral geniculate complex | Thalamus |

| | |
|---|---|
| Medial habenula | Thalamus |
| Lateral habenula | Thalamus |
| Accessory supraoptic group | Hypothalamus |
| Paraventricular hypothalamic nucleus | Hypothalamus |
| Periventricular hypothalamic nucleus, anterior part | Hypothalamus |
| Periventricular hypothalamic nucleus, intermediate part | Hypothalamus |
| Arcuate hypothalamic nucleus | Hypothalamus |
| Anterodorsal preoptic nucleus | Hypothalamus |
| Anteroventral preoptic nucleus | Hypothalamus |
| Anteroventral periventricular nucleus | Hypothalamus |
| Dorsomedial nucleus of the hypothalamus | Hypothalamus |
| Median preoptic nucleus | Hypothalamus |
| Medial preoptic area | Hypothalamus |
| Vascular organ of the lamina terminalis | Hypothalamus |
| Posterodorsal preoptic nucleus | Hypothalamus |
| Parastrial nucleus | Hypothalamus |
| Periventricular hypothalamic nucleus, posterior part | Hypothalamus |
| Periventricular hypothalamic nucleus, preoptic part | Hypothalamus |
| Subparaventricular zone | Hypothalamus |
| Suprachiasmatic nucleus | Hypothalamus |
| Ventromedial preoptic nucleus | Hypothalamus |
| Ventrolateral preoptic nucleus | Hypothalamus |
| Anterior hypothalamic nucleus | Hypothalamus |
| Lateral mammillary nucleus | Hypothalamus |
| Medial mammillary nucleus | Hypothalamus |
| Supramammillary nucleus | Hypothalamus |
| Tuberomammillary nucleus, dorsal part | Hypothalamus |
| Tuberomammillary nucleus, ventral part | Hypothalamus |
| Medial preoptic nucleus | Hypothalamus |
| Dorsal premammillary nucleus | Hypothalamus |
| Ventral premammillary nucleus | Hypothalamus |
| Paraventricular hypothalamic nucleus, descending division | Hypothalamus |
| Ventromedial hypothalamic nucleus | Hypothalamus |
| Posterior hypothalamic nucleus | Hypothalamus |
| Lateral hypothalamic area | Hypothalamus |
| Lateral preoptic area | Hypothalamus |
| Preparasubthalamic nucleus | Hypothalamus |
| Parasubthalamic nucleus | Hypothalamus |
| Perifornical nucleus | Hypothalamus |
| Retrochiasmatic area | Hypothalamus |
| Subthalamic nucleus | Hypothalamus |
| Tuberal nucleus | Hypothalamus |
| Zona incerta | Hypothalamus |
| Superior colliculus, sensory related | Midbrain |
| Inferior colliculus | Midbrain |
| Nucleus of the brachium of the inferior colliculus | Midbrain |
| Nucleus sagulum | Midbrain |
| Parabigeminal nucleus | Midbrain |
| Midbrain trigeminal nucleus | Midbrain |
| Subcommissural organ | Midbrain |
| Substantia nigra, reticular part | Midbrain |
| Ventral tegmental area | Midbrain |
| Paranigral nucleus | Midbrain |
| Midbrain reticular nucleus, retrorubral area | Midbrain |
| Midbrain reticular nucleus | Midbrain |
| Superior colliculus, motor related | Midbrain |
| Periaqueductal gray | Midbrain |
| Anterior pretectal nucleus | Midbrain |
| Medial pretectal area | Midbrain |
| Nucleus of the optic tract | Midbrain |
| Nucleus of the posterior commissure | Midbrain |
| Olivary pretectal nucleus | Midbrain |
| Posterior pretectal nucleus | Midbrain |
| Cuneiform nucleus | Midbrain |
| Red nucleus | Midbrain |
| Oculomotor nucleus | Midbrain |
| Medial accesory oculomotor nucleus | Midbrain |
| Edinger-Westphal nucleus | Midbrain |
| Trochlear nucleus | Midbrain |
| Paratrochlear nucleus | Midbrain |
| Ventral tegmental nucleus | Midbrain |

| | |
|---|---|
| Anterior tegmental nucleus | Midbrain |
| Lateral terminal nucleus of the accessory optic tract | Midbrain |
| Dorsal terminal nucleus of the accessory optic tract | Midbrain |
| Medial terminal nucleus of the accessory optic tract | Midbrain |
| Substantia nigra, compact part | Midbrain |
| Pedunculopontine nucleus | Midbrain |
| Interfascicular nucleus raphe | Midbrain |
| Interpeduncular nucleus | Midbrain |
| Rostral linear nucleus raphe | Midbrain |
| Central linear nucleus raphe | Midbrain |
| Dorsal nucleus raphe | Midbrain |
| Nucleus of the lateral lemniscus | Pons sensory |
| Principal sensory nucleus of the trigeminal | Pons sensory |
| Parabrachial nucleus | Pons sensory |
| Barrington's nucleus | Pons behavioral |
| Dorsal tegmental nucleus | Pons behavioral |
| Posterodorsal tegmental nucleus | Pons behavioral |
| Pontine central gray | Pons behavioral |
| Pontine gray | Pons behavioral |
| Pontine reticular nucleus, caudal part | Pons motor |
| Supratrigeminal nucleus | Pons sensory |
| Tegmental reticular nucleus | Pons motor |
| Motor nucleus of trigeminal | Pons motor |
| Peritrigeminal zone | Pons sensory |
| Intertrigeminal nucleus | Pons sensory |
| Superior central nucleus raphe | Pons behavioral |
| Locus ceruleus | Pons behavioral |
| Laterodorsal tegmental nucleus | Pons behavioral |
| Nucleus incertus | Pons behavioral |
| Pontine reticular nucleus | Pons motor |
| Nucleus raphe pontis | Pons behavioral |
| Subceruleus nucleus | Pons behavioral |
| Sublaterodorsal nucleus | Pons behavioral |
| Dorsal cochlear nucleus | Medulla |
| Ventral cochlear nucleus | Medulla |
| Cuneate nucleus | Medulla |
| Gracile nucleus | Medulla |
| Nucleus of the trapezoid body | Medulla |
| Nucleus of the solitary tract | Medulla |
| Spinal nucleus of the trigeminal, caudal part | Medulla |
| Spinal nucleus of the trigeminal, interpolar part | Medulla |
| Spinal nucleus of the trigeminal, oral part | Medulla |
| Paratrigeminal nucleus | Medulla |
| Abducens nucleus | Medulla |
| Facial motor nucleus | Medulla |
| Nucleus ambiguus | Medulla |
| Dorsal motor nucleus of the vagus nerve | Medulla |
| Gigantocellular reticular nucleus | Medulla |
| Inferior olivary complex | Medulla |
| Intermediate reticular nucleus | Medulla |
| Inferior salivatory nucleus | Medulla |
| Linear nucleus of the medulla | Medulla |
| Lateral reticular nucleus | Medulla |
| Magnocellular reticular nucleus | Medulla |
| Medullary reticular nucleus | Medulla |
| Medullary reticular nucleus, dorsal part | Medulla |
| Medullary reticular nucleus, ventral part | Medulla |
| Parvicellular reticular nucleus | Medulla |
| Paragigantocellular reticular nucleus, dorsal part | Medulla |
| Paragigantocellular reticular nucleus, lateral part | Medulla |
| Nucleus of Roller | Medulla |
| Nucleus prepositus | Medulla |
| Parapyramidal nucleus | Medulla |
| Lateral vestibular nucleus | Medulla |
| Medial vestibular nucleus | Medulla |
| Spinal vestibular nucleus | Medulla |
| Superior vestibular nucleus | Medulla |
| Nucleus x | Medulla |
| Hypoglossal nucleus | Medulla |
| Nucleus y | Medulla |
| Nucleus raphe magnus | Medulla |

| | |
|---|---|
| Nucleus raphe obscurus | Medulla |
| Lingula (I) | Cerebellar Cortex |
| Central lobule | Cerebellar Cortex |
| Culmen | Cerebellar Cortex |
| Declive (VI) | Cerebellar Cortex |
| Folium-tuber vermis (VII) | Cerebellar Cortex |
| Pyramus (VIII) | Cerebellar Cortex |
| Uvula (IX) | Cerebellar Cortex |
| Nodulus (X) | Cerebellar Cortex |
| Simple lobule | Cerebellar Cortex |
| Ansiform lobule | Cerebellar Cortex |
| Paramedian lobule | Cerebellar Cortex |
| Copula pyramidis | Cerebellar Cortex |
| Paraflocculus | Cerebellar Cortex |
| Flocculus | Cerebellar Cortex |
| Fastigial nucleus | Cerebellar Nuclei |
| Interposed nucleus | Cerebellar Nuclei |
| Dentate nucleus | Cerebellar Nuclei |
| Vestibulocerebellar nucleus | Cerebellar Nuclei |

***Supplementary Table S2: Region labels of our generated connectome.*** *After all modifications, the connectome used for mean-field simulations included the following regions (taken as right- and left-hemispheric, respectively). The second column shows their respective classification in terms of dynamical modeling.*

| # | Region Name | Classification for modeling |
|---|---|---|
| 1 | Frontal pole, cerebral cortex | Isocortex |
| 2 | Primary motor area | Isocortex |
| 3 | Secondary motor area | Isocortex |
| 4 | Primary somatosensory area, nose | Isocortex |
| 5 | Primary somatosensory area, barrel field | Isocortex |
| 6 | Primary somatosensory area, lower limb | Isocortex |
| 7 | Primary somatosensory area, mouth | Isocortex |
| 8 | Primary somatosensory area, upper limb | Isocortex |
| 9 | Primary somatosensory area, trunk | Isocortex |
| 10 | Primary somatosensory area, unassigned | Isocortex |
| 11 | Supplemental somatosensory area | Isocortex |
| 12 | Gustatory areas | Isocortex |
| 13 | Visceral area | Isocortex |
| 14 | Dorsal auditory area | Isocortex |
| 15 | Primary auditory area | Isocortex |
| 16 | Posterior auditory area | Isocortex |
| 17 | Ventral auditory area | Isocortex |
| 18 | Anterolateral visual area | Isocortex |
| 19 | Anteromedial visual area | Isocortex |
| 20 | Lateral visual area | Isocortex |
| 21 | Primary visual area | Isocortex |
| 22 | Posterolateral visual area | Isocortex |
| 23 | posteromedial visual area | Isocortex |
| 24 | Laterointermediate area | Isocortex |
| 25 | Postrhinal area | Isocortex |
| 26 | Anterior cingulate area, dorsal part | Isocortex |
| 27 | Anterior cingulate area, ventral part | Isocortex |
| 28 | Prelimbic area | Isocortex |

| | | |
|---|---|---|
| 29 | Infralimbic area | Isocortex |
| 30 | Orbital area, lateral part | Isocortex |
| 31 | Orbital area, medial part | Isocortex |
| 32 | Orbital area, ventrolateral part | Isocortex |
| 33 | Agranular insular area, dorsal part | Isocortex |
| 34 | Agranular insular area, posterior part | Isocortex |
| 35 | Agranular insular area, ventral part | Isocortex |
| 36 | Retrosplenial area, lateral agranular part | Isocortex |
| 37 | Retrosplenial area, dorsal part | Isocortex |
| 38 | Retrosplenial area, ventral part | Isocortex |
| 39 | Anterior area | Isocortex |
| 40 | Rostrolateral visual area | Isocortex |
| 41 | Temporal association areas | Isocortex |
| 42 | Perirhinal area | Isocortex |
| 43 | Ectorhinal area | Isocortex |
| 44 | Specific Thalamus to Frontal pole, cerebral cortex | Specific Thalamus |
| 45 | Specific Thalamus to Primary motor area | Specific Thalamus |
| 46 | Specific Thalamus to Secondary motor area | Specific Thalamus |
| 47 | Specific Thalamus to Primary somatosensory area, nose | Specific Thalamus |
| 48 | Specific Thalamus to Primary somatosensory area, barrel field | Specific Thalamus |
| 49 | Specific Thalamus to Primary somatosensory area, lower limb | Specific Thalamus |
| 50 | Specific Thalamus to Primary somatosensory area, mouth | Specific Thalamus |
| 51 | Specific Thalamus to Primary somatosensory area, upper limb | Specific Thalamus |
| 52 | Specific Thalamus to Primary somatosensory area, trunk | Specific Thalamus |
| 53 | Specific Thalamus to Primary somatosensory area, unassigned | Specific Thalamus |
| 54 | Specific Thalamus to Supplemental somatosensory area | Specific Thalamus |
| 55 | Specific Thalamus to Gustatory areas | Specific Thalamus |
| 56 | Specific Thalamus to Visceral area | Specific Thalamus |
| 57 | Specific Thalamus to Dorsal auditory area | Specific Thalamus |
| 58 | Specific Thalamus to Primary auditory area | Specific Thalamus |
| 59 | Specific Thalamus to Posterior auditory area | Specific Thalamus |
| 60 | Specific Thalamus to Ventral auditory area | Specific Thalamus |
| 61 | Specific Thalamus to Anterolateral visual area | Specific Thalamus |
| 62 | Specific Thalamus to Anteromedial visual area | Specific Thalamus |
| 63 | Specific Thalamus to Lateral visual area | Specific Thalamus |
| 64 | Specific Thalamus to Primary visual area | Specific Thalamus |
| 65 | Specific Thalamus to Posterolateral visual area | Specific Thalamus |
| 66 | Specific Thalamus to posteromedial visual area | Specific Thalamus |
| 67 | Specific Thalamus to Laterointermediate area | Specific Thalamus |
| 68 | Specific Thalamus to Postrhinal area | Specific Thalamus |
| 69 | Specific Thalamus to Anterior cingulate area, dorsal part | Specific Thalamus |
| 70 | Specific Thalamus to Anterior cingulate area, ventral part | Specific Thalamus |
| 71 | Specific Thalamus to Prelimbic area | Specific Thalamus |
| 72 | Specific Thalamus to Infralimbic area | Specific Thalamus |
| 73 | Specific Thalamus to Orbital area, lateral part | Specific Thalamus |

| | | |
|---|---|---|
| 74 | Specific Thalamus to Orbital area, medial part | Specific Thalamus |
| 75 | Specific Thalamus to Orbital area, ventrolateral part | Specific Thalamus |
| 76 | Specific Thalamus to Agranular insular area, dorsal part | Specific Thalamus |
| 77 | Specific Thalamus to Agranular insular area, posterior part | Specific Thalamus |
| 78 | Specific Thalamus to Agranular insular area, ventral part | Specific Thalamus |
| 79 | Specific Thalamus to Retrosplenial area, lateral agranular part | Specific Thalamus |
| 80 | Specific Thalamus to Retrosplenial area, dorsal part | Specific Thalamus |
| 81 | Specific Thalamus to Retrosplenial area, ventral part | Specific Thalamus |
| 82 | Specific Thalamus to Anterior area | Specific Thalamus |
| 83 | Specific Thalamus to Rostrolateral visual area | Specific Thalamus |
| 84 | Specific Thalamus to Temporal association areas | Specific Thalamus |
| 85 | Specific Thalamus to Perirhinal area | Specific Thalamus |
| 86 | Specific Thalamus to Ectorhinal area | Specific Thalamus |
| 87 | Olfactory Areas | Subcortical |
| 88 | Hippocampal Formation | Subcortical |
| 89 | Cortical Subplate | Subcortical |
| 90 | Striatum | Subcortical |
| 91 | Pallidum | Subcortical |
| 92 | Nonspecific Thalamus | Subcortical |
| 93 | Hypothalamus | Subcortical |
| 94 | Rest of Midbrain | Subcortical |
| 95 | Superior colliculus, motor related | Subcortical |
| 96 | Rest of Pons Sensory | Subcortical |
| 97 | Principal sensory nucleus of the trigeminal | Subcortical |
| 98 | Pons Motor | Subcortical |
| 99 | Pons Behavioral | Subcortical |
| 100 | Rest of Medulla | Subcortical |
| 101 | Spinal nucleus of the trigeminal | Subcortical |
| 102 | Facial motor nucleus | Subcortical |
| 103 | Inferior olivary complex | Subcortical |
| 104 | Rest of cerebellar cortex | Cerebellar Cortex |
| 105 | Ansiform lobule | Cerebellar Cortex |
| 106 | Rest of cerebellar nuclei | Cerebellar Nuclei |
| 107 | Interposed nucleus | Cerebellar Nuclei |

***Supplementary Table S3. Neural mass model parameters.*** *Parameter values of the neural mass models taken with minor modifications from (Griffiths et al., 2020), before any fitting. The gains between isocortical and thalamic populations overwrite the corresponding connectome weights.*

| Name | Unit | Value | Description |
|---|---|---|---|
| $\tau_E$ | ms | 11.11 | Excitatory population time constant |
| $\tau_I$ | ms | 11.11 | Inhibitory population time constant |
| $\tau_S$ | ms | 40 | Thalamic relay nucleus time constant |
| $\tau_{SE}$ | ms | 20 | Relay-excitatory time constant |
| $\tau_{ES}$ | ms | 20 | Excitatory-relay time constant |
| $\tau_{ER}$ | ms | 20 | Excitatory-reticular time constant |
| $\tau_{SR}$ | ms | 5 | Relay-reticular time constant |
| $\tau_{RS}$ | ms | 5 | Reticular-relay time constant |
| $w_{EE}$ | - | 1.4 | Excitatory-excitatory gain |
| $w_{EI}$ | - | 1.4 | Excitatory-inhibitory gain |
| $w_{IE}$ | - | -3.0* | Inhibitory-excitatory gain |
| $w_{II}$ | - | -0.5 | Inhibitory-inhibitory gain |
| $w_{SE}$ | - | 1.65 | Relay-excitatory gain |
| $w_{SI}$ | - | 0.2 | Relay-inhibitory gain |
| $w_{ES}$ | - | 0.6 | Excitatory-relay gain |
| $w_{ER}$ | - | 0.6 | Excitatory-reticular gain |
| $w_{RS}$ | - | -2.0 | Reticular-relay gain |
| $w_{SR}$ | - | 2.0 | Relay-reticular gain |
| $I_E$ | mV | -0.35* | Excitatory population constant input |
| $I_I$ | mV | -0.3 | Inhibitory population constant input |
| $I_S$ | mV | ** | Thalamic relay nucleus constant input |
| $I_R$ | mV | -0.8 | Thalamic reticular nucleus constant input |
| $\beta$ | - | 20.0 | Activation function gain parameter |
| $\sigma$ | - | 0.0 | Activation function threshold parameter |
| $G$ | - | *** | Global connectivity scaling factor (global coupling) |
| $w_{ij}$ | - | **** | TVMB connectome weights |
| $\tau_{ij}$ | ms | **** | TVMB connectome time delay |

** Determined according to region's connectivity indegree and fitting; the original value corresponding to region's connectivity indegree equal to zero is reported in this table (see Supplementary Note S1.1 and Methods 4.2.2 of the main manuscript).*
*** Determined according to fitting (see Supplementary Note S1).*
**** A range of values tested ({0.1, 1.0, 2.0, 3.0, 4.0, 5.0, 6.0, 7.0, 8.0, 9.0, 10.0} for fitting and final results (see Supplementary Note S1).*
***** Determined from the TVMB connectivity (see Methods 4.1 of the main manuscript)*

***Supplementary Table S4*. *Optimized neural mass model parameters*.** *This table includes values of parameter $I_S$ (thalamic relay nucleus constant input in mV) and dimensionless scaling parameters of total per region connectivity indegree, $C_{I_E}$ and $C_{W_{IE}}$, as derived from parameters $C_{st}$ (strength of scaling) and $C_{sp}$ (splitting between $C_{I_E}$ and $C_{W_{IE}}$), for different values of global connectivity scaling factor (global coupling) G, from fitting via TVMB-only simulations without any whisking-task-related parameters (all values are rounded to 3 decimal points for display in the table) (see also Supplementary Note S1.1, Supplementary Figure S9 and Methods 4.2.2 of the main manuscript).*

| $G$ | $I_S$ | $C_{st}$ | $C_{sp}$ | $C_{I_E}=C_{sp}C_{st}$ | $C_{W_{IE}}=(1-C_{sp})C_{st}$ |
|---|---|---|---|---|---|
| **0.1** | 0.095 | 1.654 | 0.188 | 0.310 | 1.344 |
| **1.0** | 0.117 | 1.181 | 0.551 | 0.651 | 0.530 |
| **2.0** | 0.116 | 1.128 | 0.531 | 0.599 | 0.529 |
| **3.0** | 0.117 | 1.029 | 0.405 | 0.416 | 0.612 |
| **4.0** | 0.106 | 0.675 | 0.516 | 0.348 | 0.327 |
| **5.0** | 0.099 | 0.852 | 0.440 | 0.375 | 0.477 |
| **6.0** | 0.103 | 0.761 | 0.447 | 0.340 | 0.421 |
| **7.0** | 0.103 | 0.797 | 0.373 | 0.298 | 0.499 |
| **8.0** | 0.096 | 0.724 | 0.376 | 0.272 | 0.452 |
| **9.0** | 0.092 | 0.616 | 0.411 | 0.253 | 0.363 |
| **10.0** | 0.085 | 0.689 | 0.364 | 0.251 | 0.439 |

***Supplementary Table S5. Values of whisking task parameters*.** *Parameters include $I_W$ (whisking nodes' constant input in mV), $G_W$ (whisking nodes' long-range coupling scaling parameter,) $g_{st}$ (scaling the strength of all pathway gains), $g_{in}$ (scaling specific to the gain of the connection from M1 to FMN), $g_{out}$ (scaling specific to the gain of the connection from CN to the specific thalamic nucleus of M1), all scaling parameters being dimensionless. The table contains also the exact scaling parameters of connections M1 to FMN and from CN to the specific thalamic nucleus of M1 following the equations $g_{FMN,M1} = g_{st} + g_{in}(99 - g_{st})$ and $g_{M1\,Spec.Thal.,CN} = g_{out}g_{st}$, respectively, whereas all other pathway gains are equal to $g_{st}$. The values are reported for different values of global connectivity scaling factor (global coupling) G, as derived from fitting via TVMB-only simulations (all values are rounded to 3 decimal points for display in the table) (see also Supplementary Note S1.1, Supplementary Figure S10 and Methods 4.1.5 of the main manuscript).*

| $G$ | **4.0** | **5.0** | **6.0** | **7.0** | **8.0** |
|---|---|---|---|---|---|
| $I_w$ | -0.362 | -0.360 | -0.342 | -0.342 | -0.371 |
| $G_w$ | 5.218 | 5.426 | 5.101 | 5.144 | 4.864 |
| $g_{st}$ | 75.839 | 75.663 | 64.788 | 63.536 | 55.766 |
| $g_{in}$ | 0.500 | 0.511 | 0.536 | 0.511 | 0.525 |
| $g_{out}$ | 0.720 | 0.625 | 0.572 | 0.498 | 0.619 |
| $g_{FMN,M1} = g_{st} + g_{in}(99 - g_{st})$ | 87.415 | 87.581 | 83.130 | 81.675 | 78.448 |
| $g_{M1\,Spec.Thal.,CN} = g_{out}g_{st}$ | 54.580 | 47.282 | 37.039 | 31.636 | 34.529 |

***Supplementary Table S6: Spiking neural network parameters.***

| ***Single neurons*** | | | | | | | | | | | | |
|---|---|---|---|---|---|---|---|---|---|---|---|---|
| | $t_{ref}$ [ms] | $C_m$ [pF] | $\tau_m$ [ms] | $E_L$ [mV] | $V_{th}$ [mV] | $V_{reset}$ [mV] | $V_{min}$ [mV] | $\lambda$; $\tau_V$ | $I_e$ [pA] | $k_{adap}$ [MH$^{-1}$] | $k_1$; $k_2$ [ms$^{-1}$] | $A_1$; $A_2$ [pA] |
| **Golgi cell** | 2.0 | 145.0 | 44.0 | -62.0 | -55.0 | -75.0 | -150.0 | 1.0; 0.4 | 16.21 | 0.22 | 0.03; 0.02 | 259.99; 178.01 |
| **Granule cell** | 1.5 | 7.0 | 24.15 | -62.0 | -41.0 | -70.0 | -150.0 | 1.0 | -0.89 | 0.02 | 0.31; 0.04 | 0.01; -0.94 |
| **Stellate / Basket cell** | 1.59 | 14.6 | 9.13 | -68.0 | -53.0 | -78.0 | - | 1.8; 1.1 | 3.71 | 2.03 | 1.89; 1.1 | 5.95; 5.86 |
| **Purkinje cell** | 0.5 | 334.0 | 47.0 | -59.0 | -43.0 | -69.0 | - | 4.0; 3.5 | 176.26 | 1.49 | 0.2; 0.04 | 157.62; 172.62 |
| **Cerebellar Nuclei (exc)** | 1.5 | 142.0 | 33.0 | -45.0 | -36.0 | -55.0 | - | 3.5; 3.0 | 75.39 | 0.41 | 0.7; 0.05 | 13.86; 3.48 |
| **Cerebellar Nuclei (inh)** | 3.0 | 56.0 | 56.0 | -40.0 | -39.0 | -55.0 | - | 0.9; 1.0 | 2.38 | 0.08 | 0.04; 0.04 | 176.36; 176.36 |

| ***Connections*** | | | |
|---|---|---|---|
| | **Weight [nS]** | **Delay [ms]** | $\tau_{syn}$ **[ms]** |
| **mossy fiber – glomerulus** | 1.0 | 1.0 | - |
| **ascending axon – Golgi cell** | 0.822 | 2.0 | 1.25 |
| **ascending axon – Purkinje cell** | 0.882 | 2.0 | 1.1 |
| **basket cell – Purkinje cell** | -0.436 | 4.0 | 2.8 |
| **basket cell – basket cell** | -0.006 | 4.0 | 2.0 |
| **glomerulus – Golgi cell** | 0.240 | 1.0 | 5.0 |
| **glomerulus – granule cell** | 0.232 | 1.0 | 1.9 |
| **Golgi cell – granule cell** | -0.148 | 2.0 | 4.5 |
| **Golgi cell – Golgi cell** | -0.00696 | 4.0 | 5.0 |
| **parallel fiber – basket cell** | 0.1 | 5.0 | 0.64 |
| **parallel fiber – Golgi cell** | 0.054 | 5.0 | 1.25 |
| **parallel fiber – Purkinje cell** | 0.136 | 5.0 | 1.1 |
| **parallel fiber – stellate cell** | 0.178 | 5.0 | 0.64 |
| **stellate cell – Purkinje cell** | -1.642 | 5.0 | 2.8 |
| **stellate cell – stellate cell** | -0.005 | 4.0 | 2.0 |
| **Purkinje cell – exc CN cell** | -0.297 | 4.0 | 0.7 |

| | | | |
|---|---|---|---|
| **mossy fiber – exc CN cell** | 0.554 | 4.0 | 1.0 |
| **Purkinje cell – inh CN cell** | -0.072 | 4.0 | 1.14 |

***Supplementary Table S7: TVMB-NEST Interface parameters.*** *Parameters include $w^{TVMB \to NEST}$ (scaling parameters of the TVMB to NEST interface; all values are rounded to 3 decimal points for display in the table), as well as $w_i^{NEST \to TVMB}$ (scaling parameters of the NEST to TVMB interface) and $b_i^{NEST \to TVMB}$ (translation parameters of the NEST to TVMB interface), where the index $i$ runs for the spiking populations mossy fibers of PrV (PrVmf), granule cells (GrC) and cerebellar nuclei (CN). All scaling parameters are dimensionless, whereas the translation ones are in units of mV/ms (all values are reported in scientific notation, rounded to three decimal places, for display in the table) (see also Supplementary Note S1.2 and Methods 4.4 of the main manuscript).*

| $G$ | **4.0** | **5.0** | **6.0** | **7.0** | **8.0** |
|---|---|---|---|---|---|
| $w^{TVMB \to NEST}$ | 46.235 | 41.737 | 42.471 | 38.808 | 37.861 |
| $w_{PrVmf}^{NEST \to TVMB}$ | 3.420e-04 | 3.332e-04 | 3.454e-04 | 3.695e-04 | 4.685e-04 |
| $w_{GrC}^{NEST \to TVMB}$ | 1.290e-05 | 1.387e-05 | 1.561e-05 | 1.676e-05 | 1.680e-05 |
| $w_{CN}^{NEST \to TVMB}$ | 2.102e-04 | 2.278e-04 | 2.342e-04 | 2.373e-04 | 2.427e-04 |
| $b_{PrVmf}^{NEST \to TVMB}$ | -5.465e-01 | -5.675e-01 | -5.885e-01 | -5.944e-01 | -6.301e-01 |
| $b_{GrC}^{NEST \to TVMB}$ | -4.165e-01 | -4.106e-01 | -4.267e-01 | -4.255e-01 | -4.242e-01 |
| $b_{CN}^{NEST \to TVMB}$ | -2.057e+00 | -2.207e+00 | -2.304e+00 | -2.336e+00 | -2.398e+00 |

***Supplementary Table S8. SNN connection lesion implementation.*** *For each lesion, the table reports the inactivated connection(s), the target population, the new value of the endogenous current for the target population, and the corresponding actual and reference rate both for the target population and the CNe. MOS: mossy fibers; CNe: excitatory neurons in the cerebellar nuclei; PC: Purkinje cells; MLI: molecular layer interneuron; BC: basket cell; SC: stellate cell.*

| **Lesion** | **Connections** | **Target populations** | **Endogenous current new [pA]** | **Target population actual rate [Hz]** | **Target population reference rate [Hz]** | **CNe actual rate [Hz]** | **DCNe reference rate [Hz]** |
|---|---|---|---|---|---|---|---|
| MOStoCN | mossy-CNe = 0 | DCNe | 130 | 72.8 | 73.25 | 72.8 | 73.25 |
| PCtoCN | PC-CNe = 0 | DCNe | 45 | 73.15 | 73.25 | 73.15 | 73.25 |
| MLItoPC | BC-PC = 0<br>SC-PC = 0 | PC | 75 | 52 | 52.5 | 73.3 | 73.25 |
| MLItoMLI | BC-BC = 0<br>SC-SC = 0 | BC, SC | -15 | 33 | 32 | 73.5 | 73.25 |

### Supplementary Note S1. Fitting model parameters and validation

#### *S1.1. Dynamical model and whisking task parameters*

Dynamical model parameter $I_S$ (constant baseline current of the excitatory relay thalamic population), and parameters $C_{I_E}$ and $C_{W_{IE}}$ (scaling parameters that control the excitation-inhibition balance relative to the connectivity indegree of each brain region) were fitted for a range of long-range coupling scaling $G$ values ($\{0.1, 1, 2, 3, 4, 5, 6, 7, 8, 9, 10\}$). The statistical model included the prior distributions $I_S{\sim}Normal(0.1, 0.1)$, $C_{st}{\sim}Uniform(0, 2)$ (i.e., the strength of the scaling), $C_{sp}{\sim}Uniform(0, 0.6)$ (i.e., a parameter splitting the scaling between $I_E$ and $W_{IE}$), and the equations $C_{I_E}$=$C_{sp}C_{st}$ and $C_{W_{IE}}$=$(1 - C_{sp})C_{st}$. The fitting target was a vector concatenating the power spectra of M1 and S1 brain regions of both hemispheres, in the range [5, 47]Hz, in steps of 1Hz, computed with linear interpolation from data points extracted from (Popa et al., 2013) using *WebPlotDigitizer* (https://automeris.io/), and normalized by the sum of powers of all frequencies. A middle range of $G$ ([4.0, 8.0]) was selected for further simulations based on visual inspection of the goodness of fit and the resulting power spectra of validation simulations (see Supplementary Table S4 and Supplementary Figure S7 for the results). This range allowed for strong enough coupling for long range interactions to be effective, while avoiding larger values of $G$, for which the excitation-inhibition balance preservation was not achieved equally well.

For the whisking task simulations, the pathway gains $g_{ij}$ and whisker node parameters ($G_W$ and $I_W$) were fitted for the selected range of $G$ values. The statistical model included the following prior distributions for each parameter: $I_W{\sim}Uniform(-0.7, 0.35)$, $G_W{\sim}Uniform(1, 9)$, $g_{st}{\sim}\ Uniform(1, 99)$ (i.e., the strength of gains), $g_{in}{\sim}\ Uniform(0.0, 1.0)$ (i.e., scaling specific to the gain of the connection from M1 to FMN), $g_{out}{\sim}\ Uniform(0.0, 1.0)$ (i.e., scaling specific to the gain of the connection from CN to the specific thalamic nucleus of M1), and the equations $g_{FMN,M1} = g_{st}$+$g_{in}(99 - g_{st})$, $g_{M1\ Spec.Thal.,CN} = g_{out}g_{st}$, and $g_{ij} = g_{st}$ for all other connections of the whisking-specific pathway undergoing strengthening. The fitting target was a measure of the normalized drop of gamma-band coherence computed as $DCOH = \langle\big(FZ(COH_{ON}) - FZ(COH_{OFF})\big)/FZ(COH_{ON})\rangle$, where $COH$ denotes the M1-S1 coherence, linearly interpolated for the gamma band frequencies in the range of [25, 60]Hz in steps of 1Hz from the data extracted from (Popa et al., 2013), the ON and OFF subscripts denote the conditions with cerebellum active or not, respectively, $FZ(.)$ denotes the Fisher Z transform, and $\langle . \rangle$ the mean across frequencies within the gamma band, generating eventually a vector of two values, one per hemisphere. See Supplementary Table S5 and Supplementary Figure S8 for the results.

For fitting all the above model parameters, the SBI Python toolbox (version 0.22.0; https://sbi.readthedocs.io/en/latest/) was used on the basis of TVMB-only simulations. Following evidence for SBI's accuracy and computational performance in a previous TVB study (Hashemi et al., 2023), the Sequential Neural Posterior Estimation – Conditional algorithm (Greenberg et al., 2019) with the default Masked Autoregressive Flow density estimator was used. We sampled from the posterior distributions of the parameters, as approximated by the neural networks, with SBI's rejection sampling algorithm. Default arguments for all SBI functions were used unless explicitly mentioned. After training, SBI was applied to sample 1,000 parameter values from the posterior distributions, the average of which was used for further simulations. Moreover, 100 of those samples were used for validation simulations, the results of which were compared against the fitting targets. The robustness of fitting results for the approximated posterior distributions was validated using the metrics of statistical mean and standard deviation of the 1,000 parameter sets sampled by SBI, as well as statistical shrinkage computed as $s = 1 - \sigma_{post}^2/\sigma_{prior}^2$, where $\sigma_{prior}$ and $\sigma_{post}$ denote the standard deviation of the prior and posterior distributions, respectively. Fitting parameters $I_S$, $C_{st}$ and $C_{sp}$ was performed along two rounds with 1,000 training parameters' samples each; the second round was aiming at increasing accuracy and used as prior distributions the posterior distributions of the first round. Fitting the parameters of the whisking-specific pathway was performed in one round with 1,000 training parameters' samples.

##### S1.2. Inter-scale interface parameters

Fitting the interface parameters $w^{TVMB \to NEST}$, $w_i^{NEST \to TVMB}$ and $b_i^{NEST \to TVMB}$ for each one of the selected $G$ values followed a three-step procedure: 1) $w^{TVMB \to NEST}$ was tuned using grid search and subsequent linear interpolation aiming for a firing rate of 4.2Hz for the granule cells of AN (Chen et al., 2017). The range of $w^{TVMB \to NEST}$ and the values of $w_i^{NEST \to TVMB}$ and $b_i^{NEST \to TVMB}$ were chosen based on preliminary co-simulations. 2) The latter two parameters were computed as

$$w_i^{NEST \to TVMB} = (\langle E_i^{TVMB}(t) \rangle_{99} - \langle E_i^{TVMB}(t) \rangle_{50}) / (\langle r_i^{NEST \to TVMB}(t) \rangle_{99} - \langle r_i^{NEST \to TVMB}(t) \rangle_{50}) \quad (12)$$

$$b_i^{NEST \to TVMB} = \langle E_i^{TVMB}(t) \rangle_{50} - w_i^{NEST \to TVMB} \langle r_i^{NEST \to TVMB}(t) \rangle_{50} \,, \quad (13)$$

where $\langle . \rangle_P$ denotes the $P$th percentile of the time series and $E_i^{TVMB}(t)$ denotes time series of the activity of the $E_i$ state variable from TVMB-only simulations. Thus, we obtained a TVMB signal of almost identical baseline and amplitude in co-simulations as in TVMB-only simulations. 3) The grid search was repeated to readjust $w^{TVMB \to NEST}$ with the tuned parameters of the NEST to TVMB interface. Finally, 10 co-simulation runs with the final set of parameters were used to confirm the target granule cell rate (see Supplementary Figure S9).

#### Supplementary Note S2. Lesions to molecular layer inhibition

Independent from the normalization of the cerebellar output, removing inhibition to PCs (*MLItoPC*) significantly decreased PC (confirming literature (Middleton et al., 2008)) and CNe gamma-band power relative to mossy fibers, while lesioning the self-inhibition in the molecular layer (*MLItoMLI*) had no significant effect (Supplementary Fig. S6A). When examining coherence between cerebellar input and output, the *MLItoPC* lesion caused decreased coherence in the gamma band but, when transferring the cerebellar output to the rest of the brain, these changes in coherence were not sufficient to affect sensory and motor cortices (Supplementary Fig. S6B-C). Self-inhibition in the molecular layer had no significant impact on gamma-band power and coherence (Supplementary Fig. S6). This result suggests that while MLI inhibition is important for whisker movements during sensory-evoked whisking (Brown et al., 2025), it might be less crucial for sensorimotor integration during free whisking. On the other hand, gamma-to-theta coherence increased inside the cerebellum with *MLItoPC* and *MLItoMLI* lesion, but decreased in M1-S1, even if – as for absolute coherence – with no statistical significance.
